# TemplateFlow: FAIR-sharing of multi-scale, multi-species brain models

**DOI:** 10.1101/2021.02.10.430678

**Authors:** Rastko Ciric, William H. Thompson, Romy Lorenz, Mathias Goncalves, Eilidh MacNicol, Christopher J. Markiewicz, Yaroslav O. Halchenko, Satrajit S. Ghosh, Krzysztof J. Gorgolewski, Russell A. Poldrack, Oscar Esteban

## Abstract

Reference anatomies of the brain and corresponding atlases play a central role in experimental neuroimaging workflows and are the foundation for reporting standardized results. The choice of such references —i.e., templates— and atlases is one relevant source of methodological variability across studies, which has recently been brought to attention as an important challenge to reproducibility in neuroscience. TemplateFlow is a publicly available framework for human and nonhuman brain models. The framework combines an open database with software for access, management, and vetting, allowing scientists to distribute their resources under FAIR —findable, accessible, interoperable, reusable— principles. TemplateFlow supports a multifaceted insight into brains across species, and enables multiverse analyses testing whether results generalize across standard references, scales, and in the long term, species, thereby contributing to increasing the reliability of neuroimaging results.

---

**B**rains are morphologically variable, exhibiting diversity in such features as overall size (***Lüders et al., 2002***), sulcal curvature (***Tosun et al., 2015***), and functional topology (***Tavor et al., 2016***; ***Mars et al., 2018***). Morphological variability manifests not only in differences between brains but also in the way that a brain changes across its lifespan, as it is remodelled by development, aging, and degenerative processes (***Courchesne et al., 2000***; ***Good et al., 2001***; ***Sowell et al., 2003***). These morphological differences often correspond with the effects of interest in neuroimaging studies and hinder direct spatial comparisons between brain maps (***Brett et al., 2002***). The substantial variability within and between individual brains necessitates a means of formalizing population-level knowledge about brain anatomy and function. Neuroscientists have answered this need by creating brain atlases as references for understanding and contextualizing morphological variability. Atlases map landmarks, features, and other knowledge about the brain as annotations that are consistent across individual brains.

The development of atlases in neuroscience has accelerated knowledge discovery and dissemination. Early endeavors, epitomized by the groundbreaking work of ***Brodmann (2006***, originally published in German in 1909) and complemented by ***Von Economo and Koskinas (2008***, originally published in German in 1925), leveraged careful scrutiny of microanatomy and cytoarchitectonic properties in small numbers of brains. Talairach’s assiduous postmortem examination of a single brain (***Talairach et al., 1957***) remarkably incorporated stereotaxy by defining three spatial reference axes over the brain that allowed anchoring neural landmarks to coordinates.

This first stereotaxic atlas saw wide use with a later, improved version (***Talairach and Tournoux, 1988***). Stereotaxy also prompted the development of the earliest surgical neuronavigation systems. ***Schurr and Merrington (1978***) developed a stereotaxic apparatus to surgically induce targeted brain lesions on cats. This work informed early sectional atlases of the rodent (***Paxinos and Watson, 1997***) and macaque (***Martin and Bowden, 2000***) brains.

On account of its implicit stereotaxy, its capacity to image the entire brain, and its non-invasive acquisition protocols, magnetic resonance imaging (MRI) has revolutionized neuroscience in general and the atlasing endeavor (***Evans et al., 2012***) in particular. In combination with software instruments’ progress to map homologous features between subjects supported by regular grids (***Avants et al., 2008***) or reconstructed anatomical surfaces (***Robinson et al., 2014***), MRI has enabled researchers to create population-average maps of a particular image modality and/or particular sample with relative ease. These maps, called “templates”, are typically created by averaging features across individuals that are representative of the population of interest to a study (***Dickie et al., 2017***). As a result, atlasing endeavours have been made contingent on templates, and have shifted away from the search for a single universal neuroanatomical pattern, instead making use of increasingly large samples with the aim of representing a population average of the distribution of morphological patterns (***Evans et al., 1993***).

Although neurotypical human adults have historically been the most comprehensively templated brains (***Evans et al., 1993***; ***Mazziotta et al., 1995, 2001***; ***Evans et al., 2012***; ***Landman et al., 2012***; ***Satterthwaite et al., 2016***), researchers have also generated templates and atlases for diseased (***Dickie et al., 2015***), infant (***Matsuzawa et al., 2001***; ***Fonov et al., 2011***; ***Shi et al., 2011***), and elderly (***Buckner et al., 2004***) human populations. Similarly, nonhuman MRI templates have seen parallel development, e.g., rodents (***Calabrese et al., 2013***; ***Szulc et al., 2015***), and primates (***Schilling et al., 2017***; ***Jung et al., 2021***).

Advancing beyond the volumetric constraints of stereotaxy, researchers of the primate neocortex have also devised standard spaces based on geometric reconstructions of the cortical surface (***Fischl et al., 1999***). This surface-based approach has the advantage of respecting the intrinsic topology of cortical folds, a development that has led to further improvements in spatial localization (***Van Essen et al., 2012***; ***Coalson et al., 2018***).

Such resources as atlases and templates, which provide standardized prior knowledge, have become an indispensable component of modern neuroimaging data workflows for two cardinal reasons. First, group inference in neuroimaging studies requires that individuals’ features are aligned into a common spatial frame of reference where their location can be called standard (***Brett et al., 2002***). Second, templates engender a stereotaxic coordinate system in which atlases can be delineated or projected. Associating atlases with template coordinates also facilitates the mapping of prior population-level knowledge about the brain into images of individual subjects’ brains (for instance, to sample and average the functional MRI signal indexed by the regions defined in an atlas; ***Yeo et al. (2011***)). Because they are integral to analytic workflows, the most widely-used templates and atlases are typically distributed along with neuroimaging software libraries. Alternatively, researchers distribute their templates through, e.g., the *NeuroImaging Tools and Resources Collaboratory* [NITRC; RRID:SCR_003430], institutional websites, or data storage systems such as FigShare [RRID:SCR_004328] or Dryad [RRID:SCR_005910]). However, using the default templates of their analyses’ software toolbox is the most common practice, as shown by ***Carp (2012b***) in a review of 241 functional MRI studies.

In sum, a number of challenges have derived from management, stewardship, distribution, reuse and reporting of templates that merit attention. In an early perspective, ***Van Essen (2002***) called for connecting templates in an aggregation of databases with “powerful and flexible options for searching, selecting, and visualizing data” and stressed the importance of resource accessibility. *TemplateFlow* provides a framework that satisfies all of the aforementioned desiderata while following the “Findability, Accessibility, Interoperability, and Reusability (FAIR) Guiding Principles” (Box 1; ***Wilkinson et al., 2016***). Following the FAIR Principles, *TemplateFlow* effectively decouples standardized spatial data from software libraries while affording processing and analysis workflows (e.g. ***Esteban et al., 2017, 2019***) with the necessary flexibility to select the most appropriate template available. *TemplateFlow* comprises a cloud-based repository of human and nonhuman imaging templates —the “*TemplateFlow Archive*”, Figure 1— paired with a Python-based library —the “*TemplateFlow Client*”— for programmatically accessing template resources (Figure 2). The resource is complemented with a “*TemplateFlow Manager*” tool to upload new or update existing resources. When adding a new template, the *Manager* initiates a peer-reviewed contribution pipeline where experts are invited to curate and vet new proposals. These software components, as well as all template resources, are version-controlled. Therefore, not only does *TemplateFlow* enable “off-the-shelf” access to templates by humans and machines, it also permits researchers to share their resources with the community. To implement several of the FAIR Principles, the *TemplateFlow Archive* features a tree-directory structure, metadata files, and data files following an organization inspired by the Brain Imaging Data Structure (BIDS; ***Gorgolewski et al., 2016***). The online documentation hub and the resource browser located at TemplateFlow.org provide further details for users.

**Figure 1.**
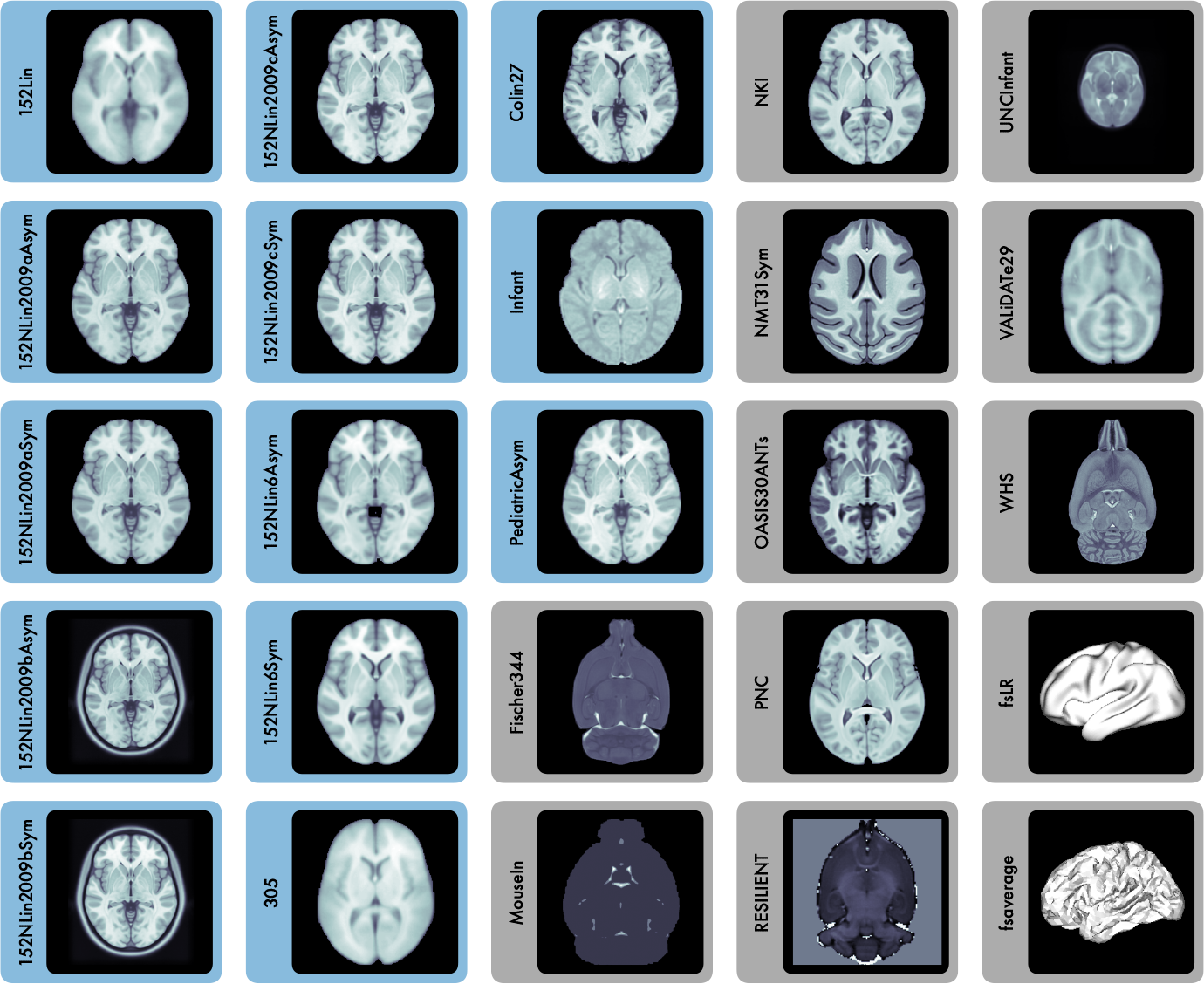
Representative views of 25 templates currently available in the *TemplateFlow Archive*. The 13 templates highlighted in blue are constituents of the Montreal Neurological Institute (MNI) portfolio. `WHS` (Waxholm space), `Fischer344`, `RESILIENT`, and `MouseIn` correspond to rodent templates. `fsaverage` and `fsLR` are surface templates; the remaining templates are volumetric. Each template is distributed with atlas labels, segmentations, and metadata files. The 25 templates displayed here are only a small fraction of those created as stereotaxic references for the neuroimaging community.

**Figure 2.**
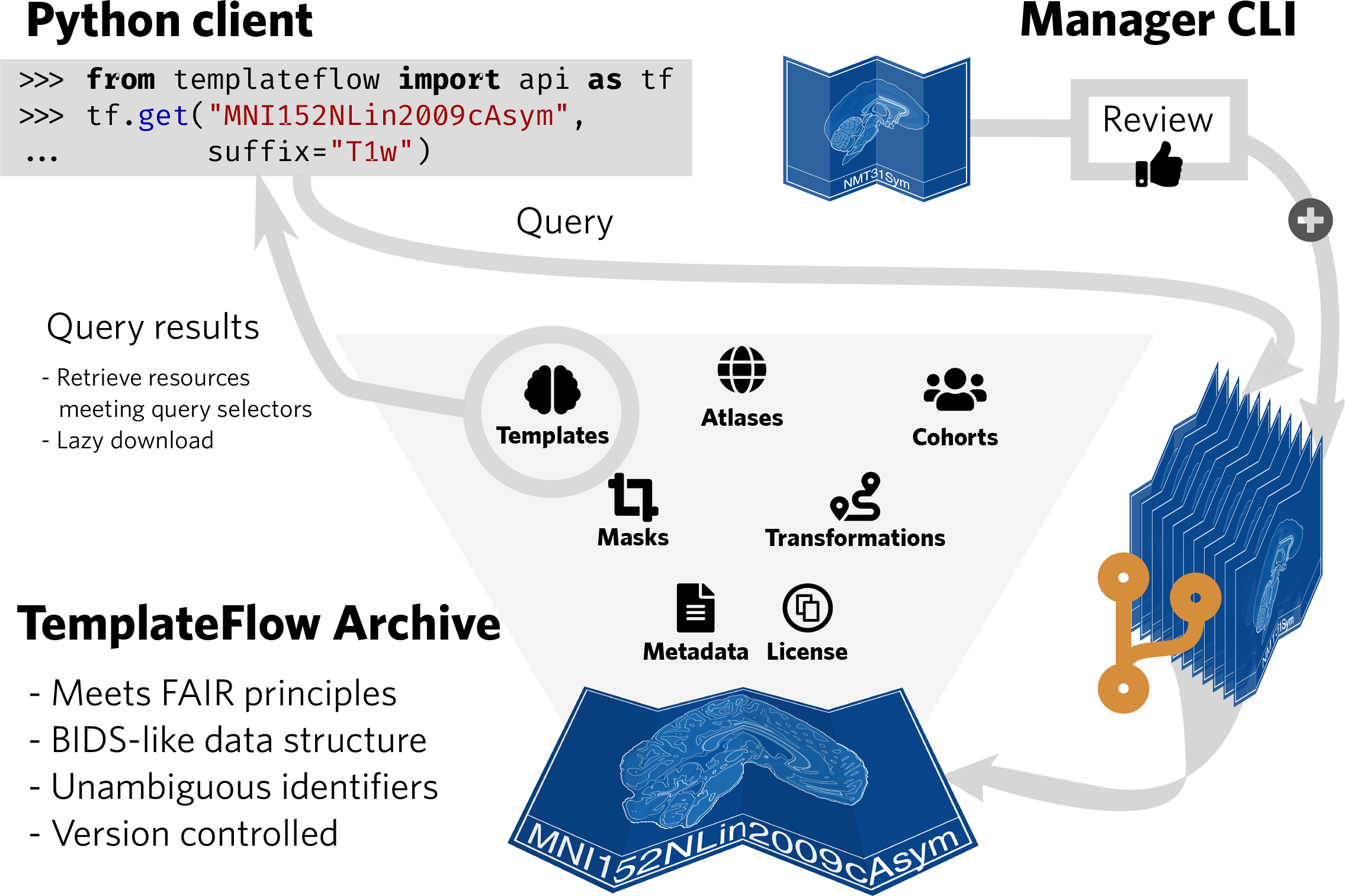
*TemplateFlow* implements the FAIR guiding principles. The *TemplateFlow Archive* can be accessed at a “low” level with *DataLad*, or at a “high” level with a *Python client*. New resources can be added through the *Manager Command Line Interface*, which initiates a peer-review process before acceptance in the *Archive*.

## Results

### BIDS-inspired structure for template and atlas archival following FAIR Principles

BIDS prescribes a file naming scheme comprising a series of key-value pairs (called “entities”). Mirroring BIDS’ patterns for participants, each template is associated with an identifier, which is an alphanumeric label. Template identifiers are unique across the *Archive*, and signified with the key `tpl`- (e.g., `tpl`-`MNI152Lin`), which is analogous to `sub`- of BIDS. Hence, every template and all associated metadata, atlases, etc. are assigned a unique and persistent identifier (Box 1, F1). The adaptation of BIDS to the domain of templates and atlases affords the tool with a robust implementation of the principles I1-3 in Box 1.

For each template, the *TemplateFlow* database includes reference volumetric template images (e.g., one T_1_- weighted MRI), a set of atlas labels and voxelwise annotations defined with reference to the template image, and additional files containing the template and atlas metadata. Correspondingly, *TemplateFlow* allows surface-based resources such as average features, geometry files, annotations, or metadata. The only requirement for feature averages and atlases sharing a unique template identifier is that they must be spatially in register, regardless of the data sampling strategy (i.e., volume, surface, or mixed, and corresponding resolution or mesh density). Although the most widely used templates generally represent MRI-derived features, *TemplateFlow* is not limited to any specific set of modalities. Table 1 has been programmatically generated by the accompanying code examples, and enumerates the templates currently distributed with the *Archive* indicating their corresponding unique identifiers and including a short description.

#### Box 1.

**The FAIR Guiding Principles**

##### To be Findable

F1. (meta)data are assigned a globally unique and persistent identifier
F2. data are described with rich metadata (defined by R1 below)
F3. metadata clearly and explicitly include the identifier of the data it describes
F4. (meta)data are registered or indexed in a searchable resource

##### To be Accessible

A1. (meta)data are retrievable by their identifier using a standardized communications protocol
  A1.1. the protocol is open, free, and universally implementable
  A1.2. the protocol allows for an authentication and authorization procedure, where necessary
A2. metadata are accessible, even when the data are no longer available

##### To be Interoperable

I1. (meta)data use a formal, accessible, shared, and broadly applicable language for knowledge representation.
I2. (meta)data use vocabularies that follow FAIR principles
I3. (meta)data include qualified references to other (meta)data

##### To be Reusable

R1. (meta)data are richly described with a plurality of accurate and relevant attributes
  R1.1. (meta)data are released with a clear and accessible data usage license
  R1.2. (meta)data are associated with detailed provenance
  R1.3. (meta)data meet domain-relevant community standards

(Reproduced from ***Wilkinson et al., 2016***, Box 2)

**Table 1.**
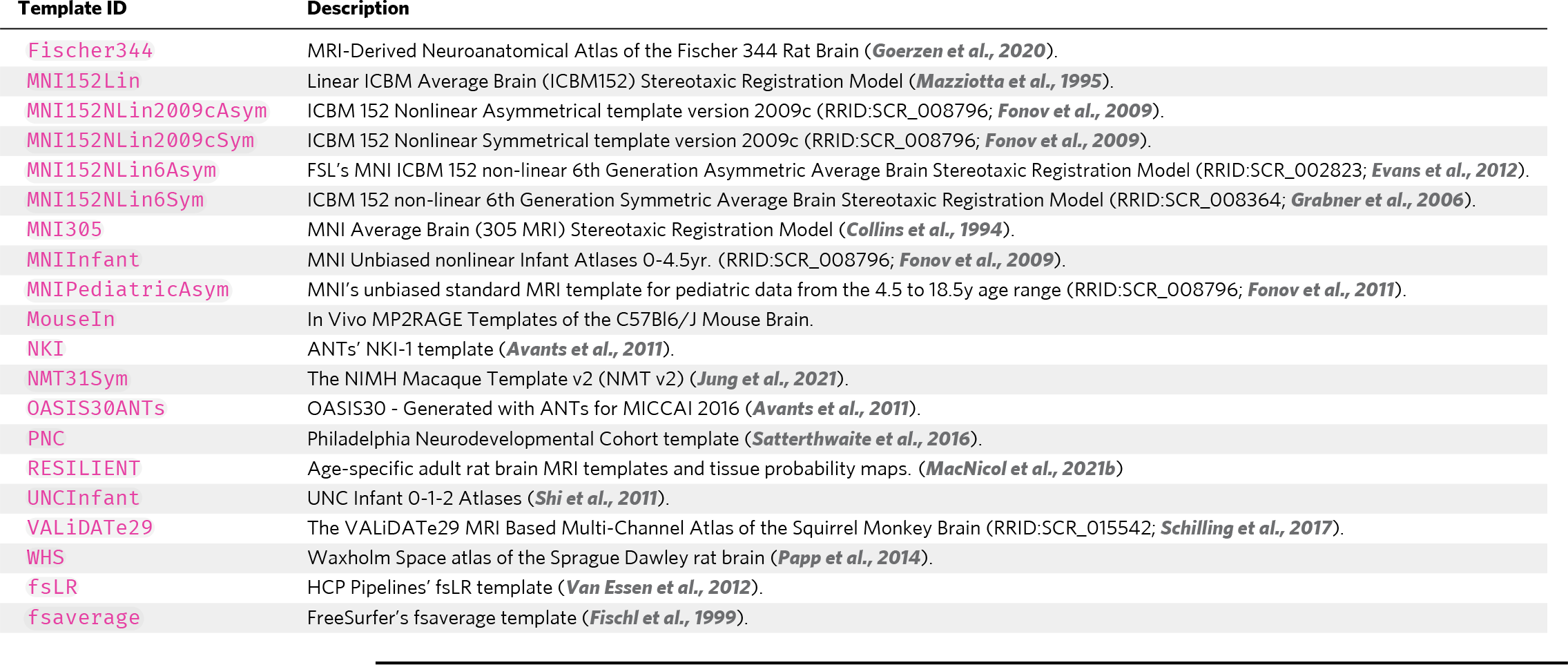
Digital templates included in *TemplateFlow*. *TemplateFlow* is designed to maximise the discoverability and accessibility of new templates, minimise redundancies in template creation, and promote standardisation of processing workflows. To enhance visibility of existing templates, *TemplateFlow* includes a web-based browser indexing all files in the *TemplateFlow Archive* (templateflow.org/browse/). This table has been automatically generated using the *TemplateFlow Client* tooling.

Template resources are described with rich metadata (Box 1, F2 and R1), ensuring that the data usage license is clear and accessible (Box 1, R1.1), data and metadata are associated with detailed provenance (Box 1, R1.2), and data and metadata follow a domain-relevant structure transferred from the neuroimaging community standards of BIDS (R1.3). Figure 3 summarizes the data types and metadata that can be stored in the *Archive*. Figure 4 provides an overview of the *Archive*’s metadata specification, showing that metadata clearly and explicitly include the identifier of the data they describe (Box 1, F3). Data and metadata are retrievable using several open, free, standard communications protocols without need for authentication (Box 1, A1) by using *DataLad* (***Halchenko et al., 2021***). Cloud storage for the *Archive* is supported by the Open Science Framework (osf.io) and Amazon’s Simple Storage Service (S3). Version control, replication, and synchronisation of template resources across filesystems is managed with *DataLad*. Leveraging *DataLad*, metadata are stored on GitHub, ensuring accessibility to metadata even when corresponding data are no longer available (Box 1, A2). *DataLad* is based on *Git* and *Git-Annex*, which index all data and metadata (Box 1, F4).

**Figure 3.**
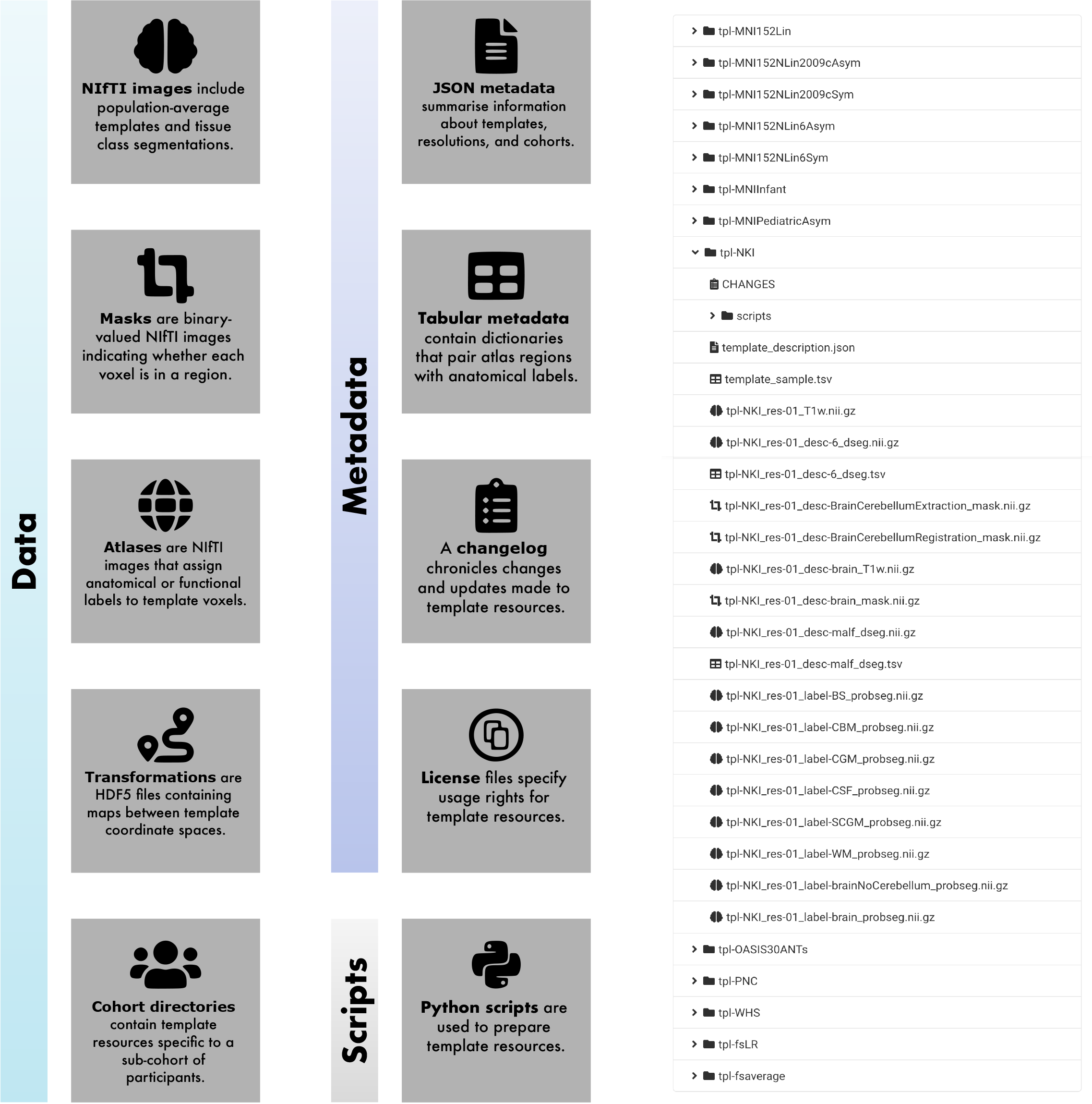
The *TemplateFlow Archive* contains template resources. Left, common file formats included in the *TemplateFlow Archive*. Right, view of the *TemplateFlow Archive*’s browser, accessible at TemplateFlow.org, with a single template resource directory expanded. Template data are archived using a BIDS-like directory structure, with top-level directories for each template. Each directory contains image files, annotations, and metadata for that template. Following BIDS specifications, volumetric data are stored in NIfTI-1 format. Further surface-based data types are supported with GIFTI (surfaces) and CIFTI-1 (mixed volumetric-and-surface data).

**Figure 4.**
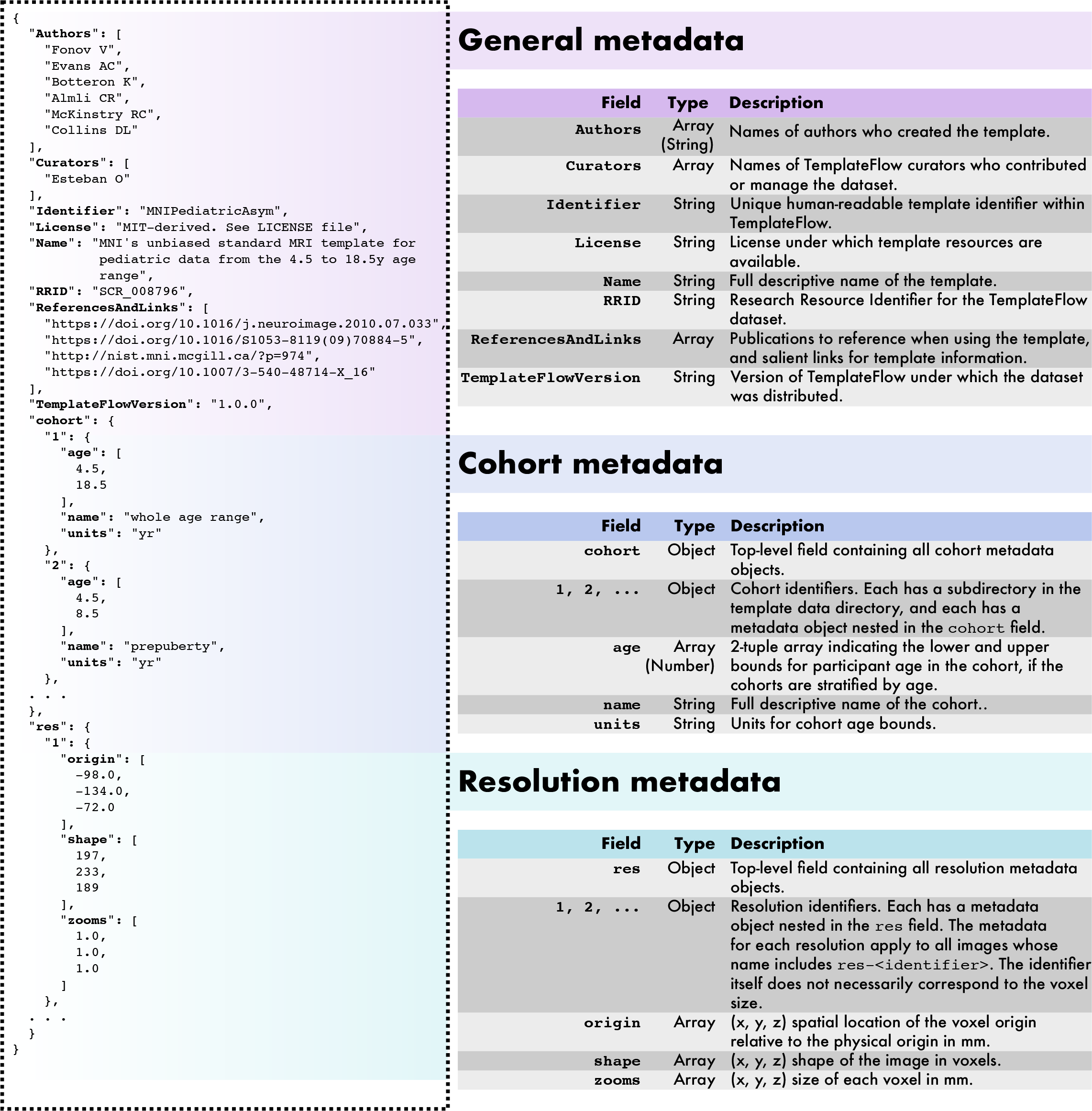
Overview of the metadata specification of the *TemplateFlow Archive*. *TemplateFlow*’s metadata are formatted as JavaScript Object Notation (JSON) files located within each template set. An example `template_description.json` metadata file is displayed at the left for `MNIPediatricAsym`. In addition to general template metadata, datasets can contain cohort-level and resolution-level metadata, which are nested within the main metadata dictionary and apply only to subsets of images in the dataset.

### *TemplateFlow* provides humans and machines with flexible and granular access to templates

*TemplateFlow*’s Python client provides human users and software tools with reliable and programmatic access to the archive. The client can be integrated seamlessly into image processing workflows to handle requests for template resources on the fly. It features an intuitive application programming interface (API) that can query the *TemplateFlow Archive* for specific files (Figure 5). The BIDS-inspired organization enables easy integration of tools and infrastructure designed for BIDS (e.g., the Python client uses *PyBIDS*; ***Yarkoni et al., 2019***). To query *TemplateFlow*, a user can submit a list of arguments corresponding to the BIDS-like key-value pairs in each entity’s file name (see Online Methods).

**Figure 5.**
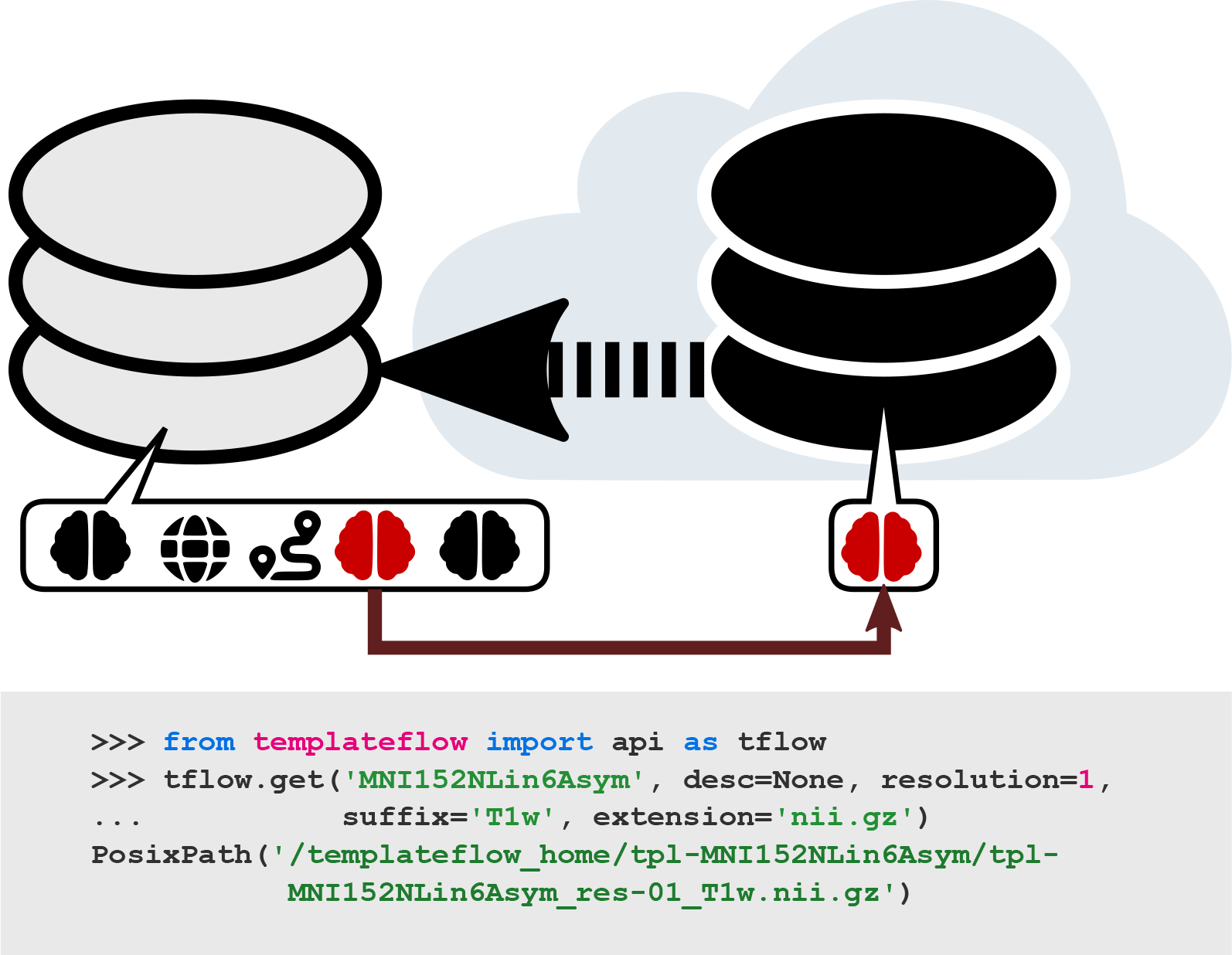
Example usage of the Python-based *TemplateFlow Client*. After importing the API, the user submits a query for the T_1_-weighted FSL version of the MNI template at 1 mm resolution. The client first filters through the archive, identifies any files that match the query, and finds their counterparts in cloud storage. It then downloads the requested files and returns their paths in the local *TemplateFlow* installation directory. Future queries for the same resource can be completed without any re-downloading.

To integrate template resources into neuroimaging workflows, traditional approaches required deploying an oftentimes voluminous tree of prepackaged data to the filesystem. By contrast, the *TemplateFlow* client implements lazy loading, which permits the base installation to be extremely lightweight. Instead of distributing neuroimaging data with the installation, *TemplateFlow* allows the user to dynamically pull from the cloud-based storage only those resources they need, as they need them. After a resource has been requested once, it remains cached in the filesystem for future utilization.

We demonstrate benefits of centralizing templates in general, and the validity of the *TemplateFlow* framework in particular, via its integration into *fMRIPrep* (***Esteban et al., 2019***), a functional MRI preprocessing tool. This integration provides *fMRIPrep* users with flexibility to spatially normalize their data to any template available in the *Archive* (see Box 2). This integration has also enabled the development of *fMRIPrep* adaptations, for instance to pediatric populations or rodent imaging (***MacNicol et al., 2021a***), using suitable templates from the archive. The uniform interface provided by the BIDS-like directory organisation and metadata enables straightforward integration of new templates into workflows equipped to use *TemplateFlow* templates. Further examples of tools leveraging *TemplateFlow* include *MRIQC* (***Esteban et al., 2017***) for quality control of MRI; *PyNets* (***Pisner and Hammonds, 2020***), a package for ensemble learning of functional and structural connectomes; *ASLPrep* (***Adebimpe et al., 2021***), an ASL pre-processing pipeline that makes use of *TemplateFlow* through *sMRIPrep* —the spin-off structural pipeline from *fMRIPrep*;— and *NetPlotBrain* (***Thompson and Fanton, 2021***), which uses *TemplateFlow* to display spatially standardized brain network data.

### *TemplateFlow* eases the dissemination of templates and opens their vetting and maintenance to the community

Beyond redistributing the templates most commonly used in the literature, the resource has the vision of becoming a centralized and standard way for researchers to disseminate their new templates. *Templateflow*’s pipeline for submission of new templates integrates peer-review with minimal technical overhead. This review process is proposed as a complement to the traditional assessment of template resources prior to publication, in which reviewers and editors focus on academic merit over accessibility and reusability potential. This submission pipeline is easily initiated with the Python-based *TemplateFlow Manager*, further described in the Online Methods.

*TemplateFlow*’s management infrastructure also eases maintenance. The lack of infrastructure for templates and atlases has given rise to static development modes where templates are packaged once and rarely revised. As a result, templates and atlases have remained outside version control because it is considered too onerous and requires informatics expertise that may exceed the resources of research teams. Although uncommon, errors in template and atlas resources have nonetheless been reported (e.g., ***Rohlfing, 2013***; ***Halchenko, 2013***), and the common denominator across these instances is template reuse for the discovery of issues. In our experience with *OpenfMRI* and *OpenNeuro* (***Markiewicz et al., 2021***), reuse is a top cause for dataset revision after the first release of data. Because of the flexibility and exposure that *TemplateFlow* affords templates and atlases will substantially lower barriers to reuse and feedback reporting, the resource will stimulate the early identification of problems and promote the request of new features to improve available assets.

Inspired by the Conda-forge community repository and the Journal of Open Source Software, the GitHub-based “templateflow” organization is a site for dialogue between members of the neuroimaging community and *TemplateFlow Archive* curators. GitHub issues offer any community member the ability to share their needs with developers and *Archive* curators, for instance by identifying templates or workflow features for potential inclusion in the project. “Pull requests” provide a means for members of the community to directly contribute code or template resources to the *TemplateFlow Archive*.

### Decoupling software and templates is indispensable for more reliable and more reproducible study designs

Although there is not yet any standard distance function that can objectively determine whether a template choice is phenotypically proximal to the study’s sample, selecting an inappropriate template may introduce so-called “template effects” that bias morphometric analyses and produce incorrect results (***Yoon et al., 2009***). For example, since most generally used templates were created with a sample of adults of European ancestry, a study involving East Asian adults might require a non-default template. More evidently, because of the relative scarcity of nonhuman imaging resources, exposure to template effects is even more pressing in the nonhuman context: e.g., is it appropriate to use a mouse template for the spatial standardization of rat images? Not only are nonhuman templates and atlases fewer, accommodation of such resources in popular software tools is generally limited. For instance, AFNI (***Cox and Hyde, 1997***) includes a rat template that can be applied in some contexts, while SPM provides functionality only through third-party add-ons (e.g., ***Sawiak et al., 2009***). Second, deviating from software defaults places a knowledge burden on the user. Once the researcher has selected a reference standard space that is suitable for their study population, if their choice is not included by default with the software they plan to use, they must then locate and download the reference template or atlas and integrate it within their analytic pipeline. To our knowledge, only AFNI have made efforts on this direction with their relatively recent @Install_<template_name> command, although the tool (i) is not designed to be compatible with other analysis alternatives, and (ii) only provides a minimal set of *TemplateFlow*’s features (i.e., resource download).

On the other hand, for tools like *TemplateFlow* that open up a wide range of options for a given study, the consequently increased methodological flexibility may become a point of concern. ***Carp (2012a***) empirically investigated the consequences of methodological flexibility in neuroimaging, demonstrating that decision points in workflows can lead to substantial variability in analysis outcomes. In a contemporaneous paper, ***Carp (2012b***) contextualized these findings vis-à-vis the inflated risk of false positives, underscoring that analytical variability degrades the reproducibility of studies only *in combination with* (intended or unintended) *selective reporting* of methods and results. Selective reporting, in this particular application, would mean that a researcher explores the results with reference to several templates or atlases and reports only those that confirm the research’s hypotheses. *TemplateFlow*’s design accounts for this risk by equipping researchers with the necessary tooling and metadata (unambiguous resource identifiers, versioning and provenance information, querying utilities, etc.) to minimize their reporting burden and ensure completeness. More recently, ***Botvinik-Nezer et al. (2020***) advocated for another approach to the problem of analytical variability: “multiverse” analyses, wherein many combinations of methodological choices are all thoroughly reported and cross-compared when presenting results. Applied to the particular choice of template and atlas combinations, it would thus be desirable to report neuroimaging results with reference to several standard spaces and determine whether the interpretations hold across those references and atlases. *TemplateFlow*’s interoperability empowers users to incorporate this type of analysis into their research by easily making template or atlas substitutions for cross-comparison. For instance, Box 2 shows how *TemplateFlow* works with *fMRIPrep* to automate preprocessing of outputs in multiple standard spaces. This facilitates assessment of the robustness of a result with respect to the template or atlas of choice in accordance with the multiverse approach. ***Botvinik-Nezer et al. (2020***) also promoted pre-registration of studies as a powerful means to fixate methodological choices before the authors have access to intermediate and final results. Here, *TemplateFlow* emerges as a powerful tool to unambiguously document template and atlas choices at pre-registration time.

## Discussion

Leveraging our experience developing an open data-sharing platform (OpenNeuro; ***Markiewicz et al., 2021***), BIDS (***Gorgolewski et al., 2016***), and related research instruments such as *MRIQC* (***Esteban et al., 2017***) and *fMRIPrep* (***Esteban et al., 2019***), we propose *TemplateFlow* as a timely resource to bridge gaps in the flexibility and reproducibility of neuroimaging studies. These gaps mostly stem from a software-bound distribution mode of templates and atlases that is the *de facto* standard practice. We argue this practice is oftentimes excessively limiting for researchers (i.e., studying the infant brain or nonhuman brain) if not problematic (i.e., template effects induced by the wrong choice of reference). More worryingly, many scientific communications assume the provenance of templates is completely specified by the choice of software library (e.g., ***Carp, 2012b***). As additional, anecdotal evidence, prompted in part by ***Carp (2012b***), in the Supplementary Materials, we show an exploration over 6,048 papers published in the specialized titles *NeuroImage* and *NeuroImage: Clinical* that contained the term “MNI” (for Montreal Neurological Institute; which has been historically established as *the* standardized space of reference). Our analysis indicates that *MNI space* can refer to any of a family of templates and is not a unique identifier. As a matter of fact, studies carried out with SPM96 (***Friston et al., 2006***) and earlier versions report their results in *MNI space* with reference to the single-subject Colin 27 average template (***Holmes et al., 1998***). However, beginning with SPM99, the software updated its definition of *MNI space* to refer to a new average of 152 subjects linearly aligned (***Mazziotta et al., 1995***). As of SPM12, different modules alternatively use the latter template or a new version created by the MNI in 2009 by means of nonlinear registration (***Fonov et al., 2011***). By contrast, the *MNI template* bundled with FSL was developed by Dr. A. Janke in collaboration with MNI researchers (***Evans et al., 2012***). Although it was generated under the guidance of and using the techniques of the 2006 release of nonlinear MNI templates, this instance is not in fact part of the official portfolio distributed by MNI. Further, FreeSurfer uses a modified version of the “MNI Average Brain (305 MRI) Stereotaxic Registration Model” (a linear MNI template based on 305 subjects; ***Evans et al., 1993***) within some early steps of the `recon`-`all` pipeline. AFNI has opted for expanding its support to all instances of the *MNI spaces* providing some of the functionalities of *TemplateFlow*. However, utilization and configuration is reserved to expert users, involved tools are not designed for compatibility, and the maintenance burden remains on the shoulders of the package developers. Issues regarding access, unequivocal identification, integration within software workflows, reuse terms, and provenance tracking are only exacerbated when the necessary template or atlas is not a software default.

We address such concerns by promoting a resource based on FAIR Principles. Indeed, it is habitual to find resources in institutional deposits such as the MNI website that lack an explicit usage license (an example at the time of writing^1^ is ***Frey et al. (2009***)). Distribution with general-purpose, manually-curated repositories such as NITRC may be more problematic. Beyond the absence of a shared data format because repositories do not mandate any standard, additional problems arise that hinder or block access and reuse. We have found templates at NITRC flagged as available for *freeware noncommercial* use as a general category, but the actual terms for reuse are not accessible. More worryingly, authors may post private resources with the appearance of being open. NITRC’s interface allows resources to be nominally released under a permissive open license, but before the data can be accessed, resources’ authors may require signing a Data Usage Agreement that forbids redistribution^2^. Further details of the advantages of *TemplateFlow* over NITRC for the particular case of template and atlas resource distribution are provided in the Table S3. Long term sustainability of the resource is ensured by using minimal cost services (all free at the time of writing) and open source tools.

## Limitations

*TemplateFlow* affords researchers substantial analytical flexibility in the choice of standard spaces of reference. Such flexibility helps researchers minimize “template effects” —by easily inserting the most adequate template— but also opens opportunities for incomplete reporting of experiments. Using *DataLad* or the *TemplateFlow Client*, researchers have at their disposal the necessary tooling for precise reporting: unique identifiers, provenance tracking, version tags, and comprehensive metadata. Therefore, the effectiveness of *TemplateFlow* to mitigate selective reporting is bounded by the user’s discretion.

As a research resource, the scope of this manuscript is limited to describing the framework and infrastructure of *TemplateFlow*, highlighting how neuroscientists can leverage this new data archive and the tooling around it. Therefore, some fundamental issues related to this work must be left for future investigation: (i) the overarching problems of cross-template and cross-atlas consistency (***Bohland et al., 2009***); (ii) comparative evaluation of methodological alternatives for producing new templates, atlases and related data; (iii) providing neuroimagers with more objective means to determine the most appropriate template and atlas choices that apply to their research, as well as better understanding “template effects”; (iv) the adequacy of original (MRI, nuclear imaging, etc.) and derived (regularly gridded images, surfaces, etc.) modalities for a specific research application; or (v) the study of the validity and reliability of inter-template registration, as well as the evaluation of such a component of the *TemplateFlow* framework. However, the proposed framework serves as an ideal keystone to investigate some of the issues above. For instance, the Supplementary Materials describe one of the resource’s tools to estimate nonlinear spatial mappings between templates that would be powerful in investigating the issue of consistency (i).

## Conclusion

We introduce an open framework for the archiving, maintenance and sharing of neuroimaging templates and atlases called *TemplateFlow* that is implemented under FAIR data sharing principles. We describe the current need for this resource in the domain of neuroimaging, and further discuss the implications of the increased analytical flexibility this tool affords. These two facets of reproducibility —availability (under FAIR guiding principles) of prior knowledge required by the research workflow, and the analytical flexibility such availability affords— are ubiquitous concerns across disciplines. *TemplateFlow*’s approach to addressing both establishes a pattern broadly transferable beyond neuroimaging. We envision *TemplateFlow* as a core research tool undergirding multiverse analyses —assessing whether neuroimaging results are robust across population-wide spatial references— as well as a stepping stone towards the quest of mapping anatomy and function across species.

## Acknowledgments

The development of this resource was supported by the Laura and John Arnold Foundation (RAP and KJG), the NIBIB (R01EB020740, SSG; 1P41EB019936-01A1 SSG, YOH), NIMH (RF1MH121867 RAP, OE; R24MH114705 and R24MH117179, RAP; 1RF1MH121885 SSG), NINDS (U01NS103780, RAP), and NSF (CRCNS 1912266, YOH). RL is funded by the Wellcome Trust (209139/Z/17/Z). EM was supported by the UK Medical Research Council (MR/N013700/1) and King’s College London. OE acknowledges financial support from the SNSF Ambizione project “Uncovering the interplay of structure, function, and dynamics of brain connectivity using MRI” (grant number PZ00P2_185872).

## Ethical compliance

We complied with all relevant ethical regulations. This resource reused publicly available data derived from studies acquired at many different institutions. Protocols for all of the original studies were approved by the corresponding ethical boards.

## Code & data availability statement

All the software components discussed in this paper are available under the Apache 2.0 license, accessible as reposi- tories of https://github.com/templateflow. All templates and associated data are available under corresponding open licenses and accessible as described in the manuscript.

## Author contributions

Conceptualization: RC, CJM, KJG, RAP, OE Data curation: RC, EM, OE Topics analysis: RC, RL, OE Funding acquisition: KJG, RAP, OE Methodology: RC, OE Project administration: RAP, OE Resources: YOH, SSG, KJG, RAP, OE Software & Documentation: RC, WHT, MG, EM, CJM, YOH, OE Supervision: RAP, OE Validation: RC, WHT, EM, OE Visualization: RC, RL. Writing – original draft: RC & OE Writing – review & editing: RC, RL, WHT, MG, EM, CJM, YOH, SSG, KJG, RAP, OE.

## Methods

*TemplateFlow* comprises four cardinal components: (i) a cloud-based archive, (ii) a Python client for program- matically querying the archive, (iii) automated systems for synchronizing and updating archive data, and (iv) inter-template registration workflows. Here, we discuss the details of each component’s implementation in turn, as well as the manner in which they interact with one another to form a cohesive whole.

### The *TemplateFlow Archive*

The archive itself comprises directories of template data in cloud storage. For redundancy, the data are stored on both Google Cloud using the Open Science Framework (OSF) and on Amazon’s Simple Storage Service (S3). Prior to storage, all template data must be named and organized in directories conforming to a data structure adapted from the Brain Imaging Data Structure (BIDS) standard (***Gorgolewski et al., 2016***). The precise implementation of this data structure is a living document and is detailed on the *TemplateFlow* homepage (http://www.templateflow.org). We detail several critical features here.

The archive is organized hierarchically, and descriptive metadata follow a principle of inheritance: any metadata that apply to a particular level of the archive also apply to all deeper levels (Figure 3). At the top level of the hierarchy are directories corresponding to each archived template. If applicable, within each template directory are directories corresponding to sub-cohort templates. Names of directories and resource files constitute a hierarchically ordered series of key-value pairs terminated by a suffix denoting the datatype. For instance, `tpl`-`MNIPedi- atricAsym_cohort`-`3_res`-`high_T1w.nii.gz` denotes a T_1_-weighted (T1w) template image file for resolution “high” of cohort “3” in the “MNIPediatricAsym” template (where the definitions of each resolution and cohort are specified in the template metadata file, *TemplateFlow Archive*). The most common *TemplateFlow* datatypes are indexed in Table S1; an exhaustive list is available in the most current version of the BIDS standard (https://bids.neuroimaging.io/).

Within each directory, template resources include image data, atlas and template metadata, transform files, licenses, and curation scripts. All image data are stored in gzipped NIfTI-1 format and are conformed to RAS+ orientation (i.e., left-to-right, posterior-to-anterior, inferior-to-superior, with the affine qform and sform matrices corresponding to a cardinal basis scaled to the resolution of the image). Template metadata are stored in a JavaScript Object Notation (JSON) file called `template_description.json` ; an overview of metadata specifications is provided in Figure 4. In brief, template metadata files contain general template metadata (e.g., authors and curators, references), cohort-specific metadata (e.g., ages of subjects included in each cohort), and resolution-specific metadata (e.g., dimensions of images associated with each resolution). Atlas metadata are often stored in TSV format and specify the region name corresponding to each atlas label. Transform files are stored in HDF5 format and are generated as a diffeomorphic composition of ITK-formatted transforms mapping between each pair of templates.

The archive has a number of client-facing access points to facilitate browsing of resources. Key among these is the archive browser on the *TemplateFlow* homepage, which indexes all archived resources and provides a means for researchers to take inventory of possible templates to use for their study.

### The Python client

*TemplateFlow* is distributed with a Python client that can submit queries to the archive and download any resources as they are requested by a user or program. Valid query options correspond approximately to BIDS key-value pairs and datatypes. A compendium of common query arguments is provided in Table S1, and comprehensive documentation is available on the *TemplateFlow* homepage.

When a query is submitted to the *TemplateFlow* client, the client begins by identifying any files in the archive that match the query. To do so, it uses *PyBIDS* (***Yarkoni et al., 2019***), which exploits the BIDS-like architecture of the *TemplateFlow Archive* to efficiently scan all directories and filter any matching files. Next, the client assesses whether queried files exist as data in local storage. When a user locally installs *TemplateFlow*, the local installation initially contains only lightweight pointers to files in OSF cloud storage. These pointers are implemented using *DataLad* (***Halchenko et al., 2021***), a data management tool that extends git and git-annex. *TemplateFlow* uses *DataLad* principally to synchronize datasets across machines and to perform version control by tracking updates made to a dataset.

If the queried files are not yet synchronized locally (i.e., they exist only as pointers to their counterparts in the cloud), the client instructs *DataLad* to retrieve them from cloud storage. In the event that *DataLad* fails or returns an error, the client falls back on redundancy in storage and downloads the file directly from Amazon’s S3. When the client is next queried for the same file, it will detect that the file has already been cached in the local filesystem. The use of resource pointers with the client thus enables lazy loading of template resources. Finally, the client confirms that the file has been downloaded successfully. If the client detects a successful download, it returns the result of the query; in the event that it detects a synchronization failure, it displays a warning for each queried file that encountered a failure.

Continued functionality and operability of the client is ensured through an emphasis on maximizing code coverage with unit tests. Updating the client requires successful completion of all unit tests, which are automatically executed by continuous integration (CI) and continuous delivery (CD) services connected to GitHub. CI and CD also keep the web-based archive browser up to date by automatically indexing data files.

### Ancillary and managerial systems

*TemplateFlow* includes a number of additional systems and programs that serve to automate stages of the archive update process, for instance addition of a new template or revision of current template resources. To facilitate the update and extension process, *TemplateFlow* uses GitHub actions to automatically synchronize dataset information so that all references remain up-to-date with the current dataset. These actions are triggered whenever a pull request to *TemplateFlow* is accepted. For example, GitHub actions are used to update the browser of the *TemplateFlow Archive* so that it displays all template resources as they are uploaded to the archive.

Whereas the *TemplateFlow Client* synchronizes data from cloud storage to the local filesystem, a complementary *TemplateFlow Manager* handles the automated synchronization of data from the local filesystem to cloud storage. The Python-based manager is also used for template intake, i.e., to propose the addition of new templates to the archive. To propose adding a new template, a user first runs the *TemplateFlow Manager* using the `tfmgr add <template_id>` --`osf`-`project <project_id>` command.

The manager begins by using the *TemplateFlow* client to query the archive and verify that the proposed template does not already exist. After verifying that the proposed template is new, the manager synchronizes all specified template resources to OSF cloud storage. It then creates a fork of the `tpl`-`intake` branch of the *TemplateFlow* GitHub repository and generates an intake file in Tom’s Obvious Minimal Language (TOML) markup format; this intake file contains a reference to the OSF project where the manager has stored template resources. The *TemplateFlow Manager* commits the TOML intake file to the fork and pushes to the user’s GitHub account. Finally, it retrieves template metadata from `template_description.json` and uses the metadata to compose a pull request on the `tpl`-`intake` branch. This pull request provides a venue for discussion and vetting of the proposed addition of a new template.

#### Box 2.

**Integration of *TemplateFlow* in processing workflows**

*TemplateFlow* maximizes the accessibility and reuse potential of templates and atlases. For example, let’s reuse the base configuration file for FSL FEAT we proposed in our paper (***Esteban et al., 2019***). The design file `design.fsf` specifies a simple preprocessing workflow with FSL tools. The simplified code listing below shows that, just to make non-default templates available to FSL using the graphical user interface (GUI), at least five steps are necessary:

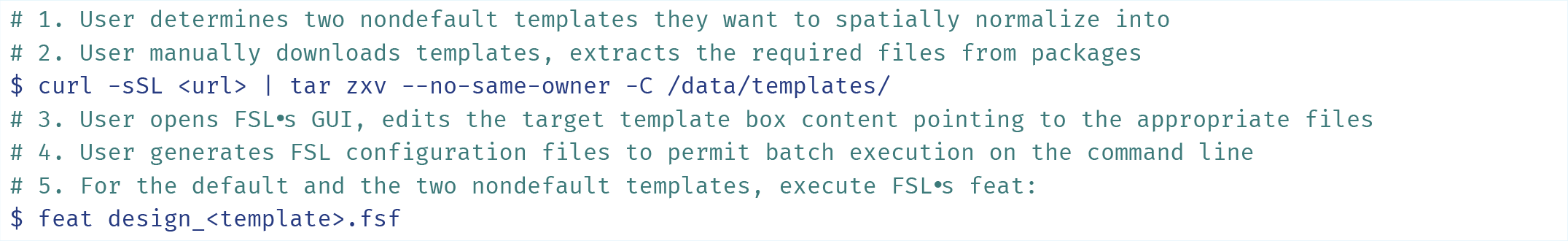

The outputs of each `feat design_<template>.fsf` call will follow the pre-specified patterns of FSL, with whatever customization the user has introduced into the design file. The user, therefore, must then adapt the downstream analysis tools to correctly interpret the derived dataset, in each standard space, or reformat the output dataset according to the expectations of the analysis tools.

The user is also responsible for all aspects of provenance tracking and adequately reporting them in their communications. Information such as version of the template (or download date), citations to relevant papers, and other metadata (e.g., RRIDs) must be accounted for manually throughout the research process.

In contrast, tools using *TemplateFlow* dramatically simplify the whole process (note that `MNI152NLin2009cAsym` and `OASIS30Ants` are the two templates not found within the FSL distribution, and `MNI152NLin6Asym` denotes *FSL’s MNI space* (i.e., the default FSL template):

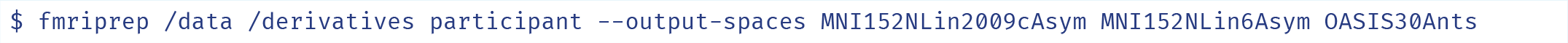

*fMRIPrep* generates the results with BIDS-Derivatives organization for the three templates. The tool also leverages *TemplateFlow* to generate a *boilerplate citation text* that includes the full names, versions and references to credit the template’s authors for each of the templates involved. *fMRIPrep* internally stages one spatial normalization workflow for each of the output spaces. Each of these normalization sub-workflows uses a simple line of Python code to retrieve the necessary resources from *TemplateFlow* using the *TemplateFlow Client* interface (Figure 5):

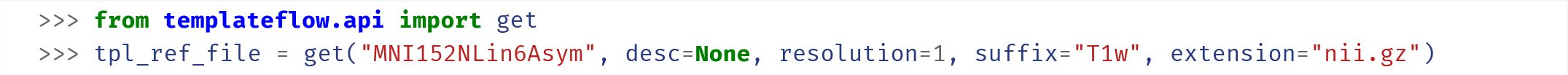

One detail overseen in the FSL example is that, for a robust spatial normalization process, a precise binary mask of the brain is generally used. While FSL would require the user to manually set this mask up in the GUI, in the case of *TemplateFlow*, it requires a second minimal call:

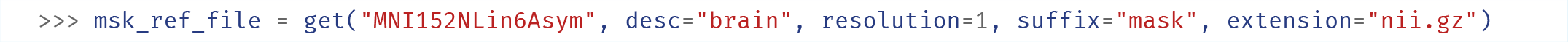

These examples are extreme simplifications of what a pipeline developer can automate and make more robust by integrating *TemplateFlow* in their workflows.

For further examples on how *TemplateFlow* can be leveraged, *PyNets* is a package for ensemble learning of functional and structural connectomes (***Pisner and Hammonds, 2020***), and *NetPlotBrain* for visualization (***Thompson and Fanton, 2021***).

**Figure 6.**
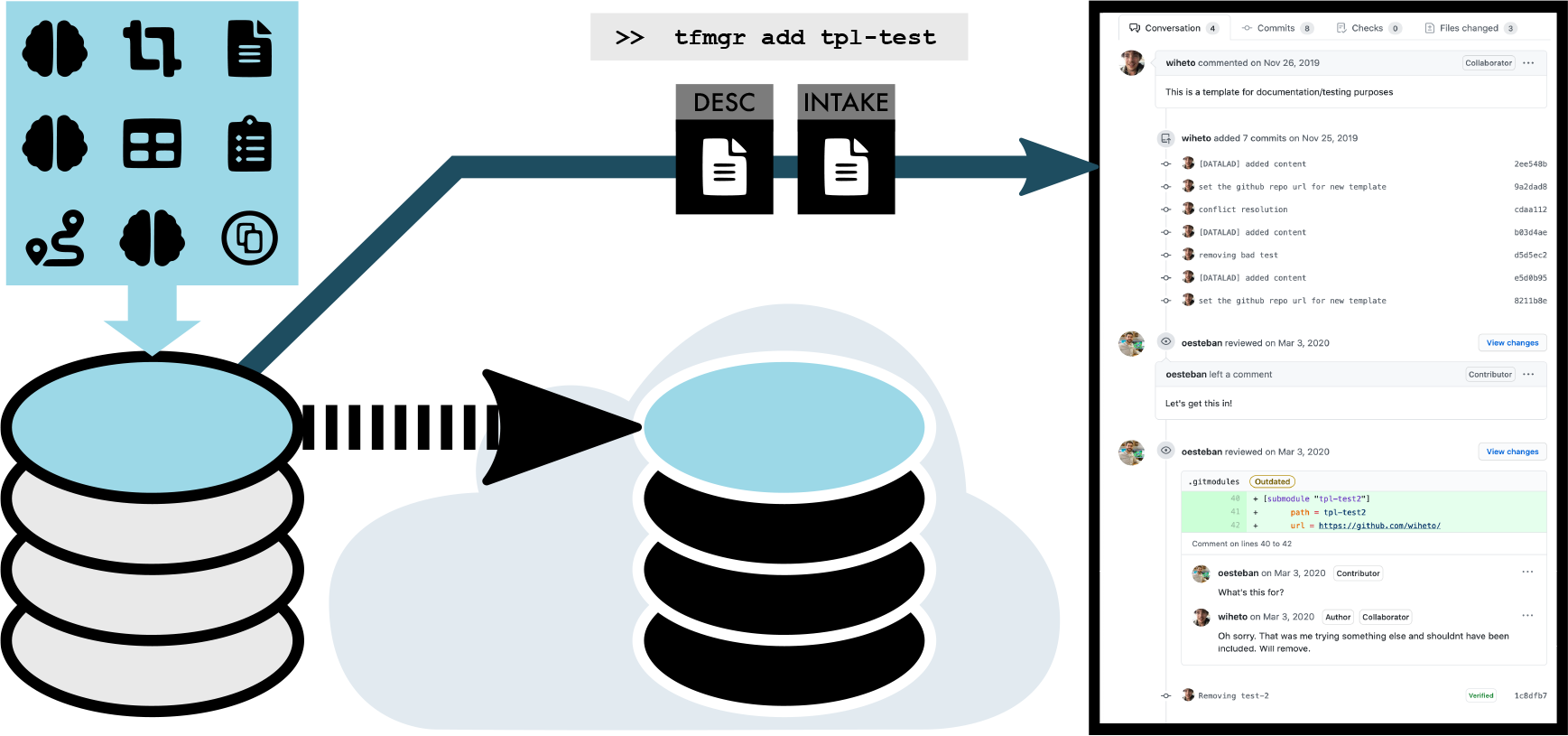
Contributing a new resource with the *TemplateFlow Manager*. To contribute a new template to *TemplateFlow*, members of the community first organize template resources to conform to the BIDS-like *TemplateFlow* structure. Next, `tfmgr` (the *TemplateFlow Manager*, see Table S2) synchronizes the resources to OSF cloud storage and opens a new pull request proposing the addition of the new template. A subsequent peer-review process ensures that all data are conformant with the *TemplateFlow* standard. Finally, *TemplateFlow* curators merge the pull request, thereby adding the template into the archive.

## Supplementary Materials

### Inter-template registration workflow

To enable the flow of knowledge across template spaces, *TemplateFlow* includes a workflow for computing robust transformations between any pair of adult human template spaces. To compute a transformation between two template spaces, the inter-template registration workflow makes use of 10 of the high-quality T_1_-weighted (T1w) adult human brain images used in the creation of the MNI 152 template portfolio. In the first step of the workflow, these 10 images are registered to both template spaces using the symmetric normalization (SyN) algorithm (***Avants et al., 2008***). SyN is an algorithm for nonlinear registration of images, which has the objective of identifying a diffeomorphic transformation that maximizes the cross-correlation between the (*moving*) image it transforms and a target (*fixed* or *reference*) image. Here, each of the 10 subject T1w images is input as a moving image, and each of the two template spaces to be connected is input as a reference image to a single transform. Thus, 20 total transforms are estimated in the first stage of the workflow. We use the SyN algorithm in conjunction with an affine initialization, which computes a preliminary alignment between images before they are input to the SyN algorithm.

Next, in the second stage of the workflow, we exploit the multi-channel registration capacity of the SyN algorithm to estimate a single, composite template-to-template transformation in each direction. In further detail, the second stage begins with a brain extraction step, which uses template-based priors to remove skull and extracranial voxels from each of the 10 T1W images aligned to each template. Next, we correct for intensity non-uniformities (INU) in each image using the N4ITK bias field correction algorithm (***Tustison et al., 2010***). The SyN algorithm then permits each of the skull-stripped, INU-corrected images to be input as registration targets on a separate registration channel. Thus, we optimize a 10-channel registration, wherein the 10 moving images are the 10 skull-stripped subject T1w images that we registered to one template space in the first step, and the 10 reference images are the same skull-stripped T1w images, this time registered to the other template space. To estimate the inverse template-to-template transform, we repeat this process, reversing the assignments of reference and moving images. Thus, the workflow computes a single transformation that simultaneously optimizes the alignment between all 10 images in both coordinate spaces.

If *FreeSurfer*-formatted `.aparc` parcellation files are available for each image, they are warped to each template space. The warped parcellation labels are then used as inputs to a joint label fusion (JLF) procedure (***Wang et al., 2013***), which returns both a deterministic parcellation and a set of label prior maps for each template. Additionally, reconstructed subject surfaces are warped into each template space using a point-wise version of the transformations computed in the first step, applied to the vertices of each subject’s cortical surface. After vertices are transformed, surfaces are reconstituted using Poisson surface reconstruction (***Kazhdan and Hoppe, 2013***).

### “MNI space” text mining analysis

To investigate the coupling between software libraries and standard spaces, we conducted an exploratory text mining analysis of the term “MNI” in the neuroimaging literature. For this, we used the Elsevier API to download the entire corpus of two leading journals of neuroimaging methodology, *NeuroImage* and *NeuroImage: Clinical*. In this way, we retrieved a total 16,812 full-text articles that were subsequently segmented into lists of sentences. A scan of these sentences revealed 14,870 sentences across 6,048 articles that contained the word “MNI”. Sentences were cleaned (i.e., removing punctuation, single letters, accents, numbers) and tokenized into words, which were subsequently lemmatized (i.e., converted to base form) using the NLTK wordnet lemmatizer. From the lemmatized words, we filtered out stopwords (i.e., NLTK stopwords and a custom list) and included words with a frequency above 10 as part of our “dictionary”; this yielded a dictionary size of 2,324 words.

Next, we computed a sparse dictionary size by article count matrix (i.e., 2,324 × 6,048), on which we performed topic modelling with latent Dirichlet allocation (LDA; ***Blei et al., 2003***, implementation from scikit-learn with the learning decay hyperparameter set to 0.7). The number of topics (*k*=15) was selected by identifying the LDA model yielding the lowest perplexity (***Blei et al., 2003***) sweeping the interval [8-16] for the parameter. The 20 words from the dictionary that loaded the highest on the 15 topics were visualized using word clouds. Topics were sorted by descending dominance, and the dominance fraction (number of articles where the topic is the most loaded with respect to the total 6,048 documents) was included above the corresponding topic’s word cloud (Figure S1).

Our topic model identified a clear coupling between the reporting of spatial standardization and software libraries. Out of 15 topics we modeled, two of the most dominant topics (Topics 3 and 5) contained software tool names as well as the names of related scientists. As shown in S1, around 500 articles (each term) contained either “SPM” (9% of the documents, Topic #3) or “FSL” (8%, Topic #5) in the same sentence as “MNI”. Interestingly, the two words do not ever appear together, suggesting that researchers stick with one or another in their analyses. Additional topics that seemingly relate to the provenance of templates and atlases–beyond the ubiquitous “Montreal”, “Neurological” and “Institute” for MNI–are those that ranked #4, #13, #14, which include “SPM” and other terms such as “McGill”, “Wellcome”, “UCL”, or the “statistical”, “parametric”, and “mapping” in SPM (see S1). The remainder of topics appears to relate to miscellaneous aspects of spatial standardization, such as “anatomical”, “smoothness”, “standard”, “normalization”, “(re)align/alignment”, etc., with no immediately apparent relationship to the origin of the resource. Nonetheless, our results suggest that the MNI templates bundled with SPM and FSL have historically gained broader currency as a result of the widespread use of these software libraries.

Our text mining results are consistent with a previous investigation by ***Carp (2012b***) in the domain of functional MRI, which similarly illustrated a strong coupling between software library and authors’ choices of templates and atlases. In this prior work, ***Carp (2012b***) analysed 241 functional MRI studies, 90.9% of which reported normalizing brain images to a common template. Of those, 79.0% indicated the target space used for spatial normalization. Few studies reported critical parameters such as image modality, and only 50 out of 241 specified the template image, out of which 26.0% used “the MNI152 template”, and 26.0% the “SPM library’s echo-planar imaging template”. Unfortunately, template selection is seen as a default parameter of the software library, which leads to the assumption that the target normalization space is implicitly reported by identifying the software tools of choice. The risks and limitations of this reporting scheme are further underscored by a recent comparison of analytic outcomes across software libraries. In this study, ***Bowring et al. (2019***) implemented analogous image processing pipelines using tools from each of three software suites (AFNI, FSL and SPM) in order to identify challenges to reproducing published studies with openly shared raw data. When discussing the differences among software pipelines, they noted that, “while all packages are purportedly using the same MNI atlas space, an appreciable amount of activation detected by AFNI and FSL fell outside of SPM’s analysis mask.”

Furthermore the coupling between templates and software likely limits the use of templates other than those defaulted by the software. Custom templates (i.e., those not included as a default option for the software tool) range from population-specific templates to *ad hoc* templates created by averaging images of the study at hand. In some settings, the use of default templates risks introducing “template effects” that confound the interpretation of results (such as those introduced when an adult template is ised in pediatric imaging studies, ***Yoon et al., 2009***). As the target population moves away from the population used to create a default template —generally, a neurotypical adult population— “template effects” become more concerning and custom templates more necessary. The problem is exacerbated in the case of nonhuman imaging, as the scarcity (or absence) of specific templates available within software packages hinders already challenging translational endeavors. Further, the consistency across templates and atlases is reportedly low (***Bohland et al., 2009***), and although there has not been any programmatic comparison to understand the extent to which this inconsistency alters the spatial interpretation of results, it is reasonable that templates and atlases introduce a decision point and therefore are sources of some analytical variability.

**Figure S1.**
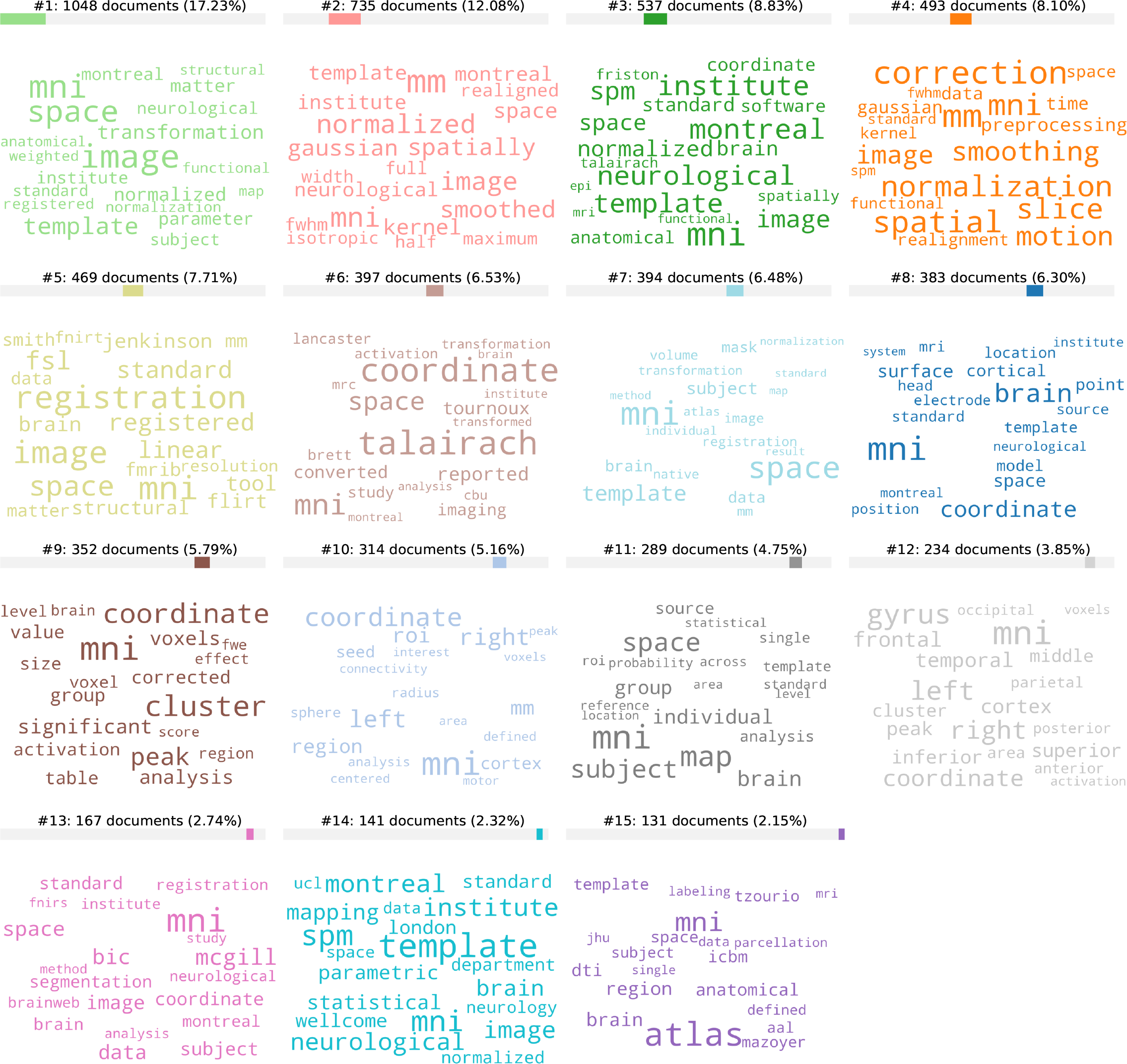
The FSL and SPM software tools associate with dominant topics of sentences including the term “MNI” across the literature. We performed topic modeling with latent Dirichlet allocation (LDA; ***Blei et al., 2003***) on text sentences extracted from 6,048 articles that contained the word “MNI”. For each topic identified, the 20 words with the highest loadings on that topic are displayed in a word cloud with larger font size indicating higher loading of the word on the corresponding topic. Word clouds are sorted by descending topic dominance. Ranking and relative dominance are shown above each topic’s cloud.. Two top-dominant topics —#3 and #5— are associated with SPM and FSL respectively.

#### Supp. Box S1.

**Quick start with the *TemplateFlow Client* API**

##### Finding templates

At the time of writing there are 25 templates available within the resource, and all the unique identifiers can be accessed with the `templates()` method:

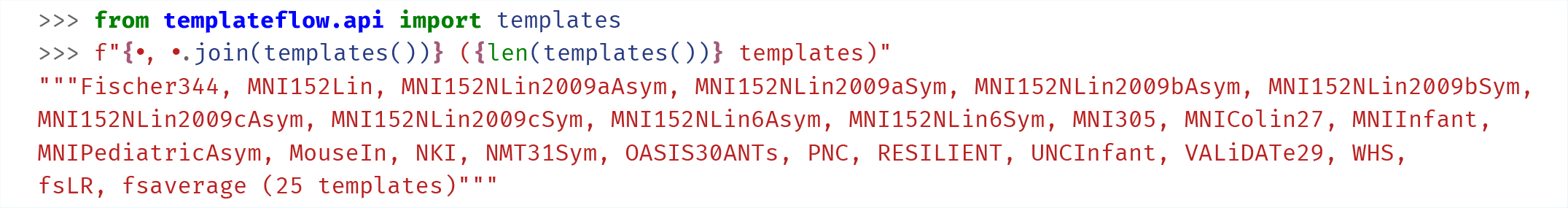

##### Accessing Metadata

We can query metadata associated to individual data files (e.g., a volume or a surface) or general metadata of the template. For example, the `get_metadata(<template_id>)` returns the general metadata as a dictionary. Hence, consulting the full name corresponding to some template identifiers yields:

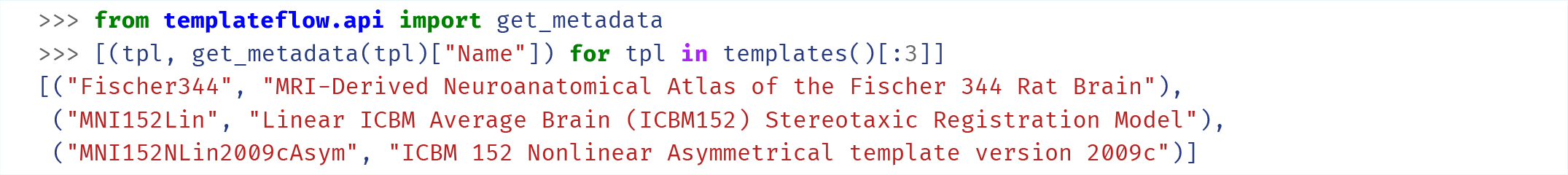

Similarly, we can check the license of a given template:

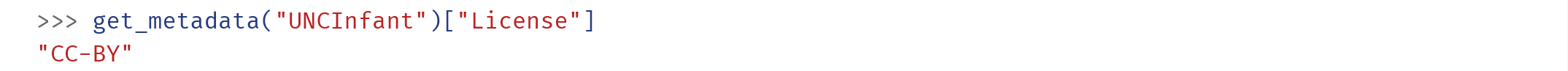

Or the proper citations (please note that the output in this example has been manipulated for demonstration purposes):

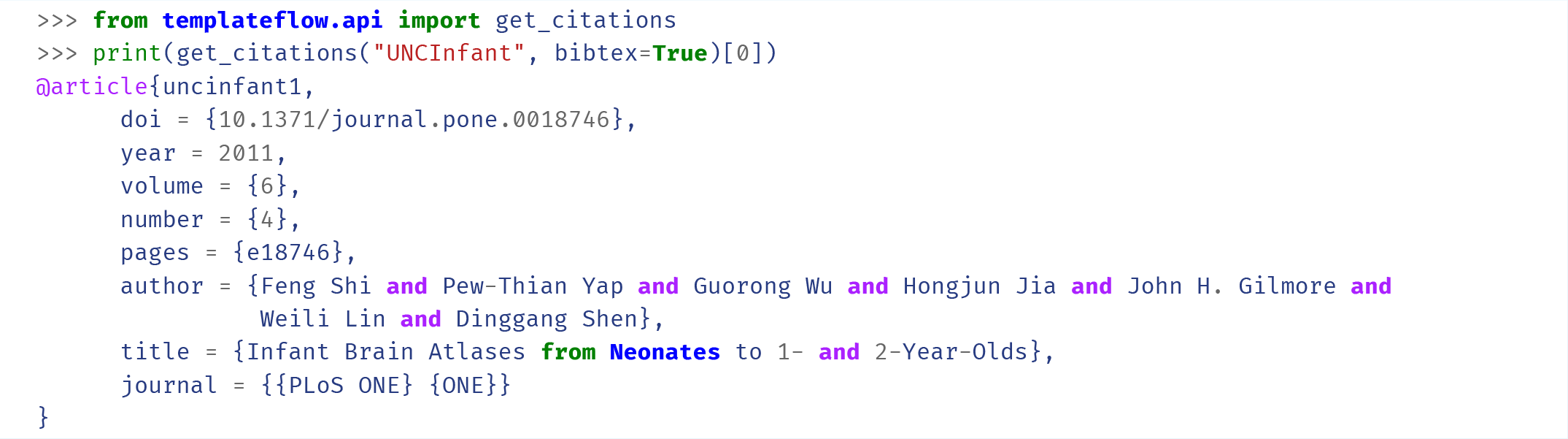

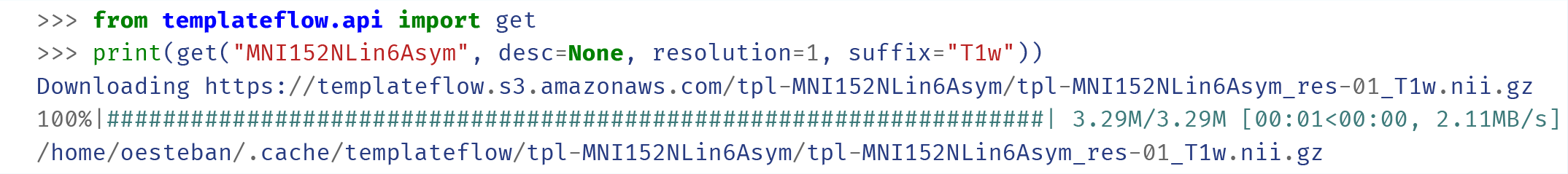

**Table S1.** *TemplateFlow* data entities.

| Data entity | API query example | Description |
| --- | --- | --- |
| Template | "MNI152Lin" | The template dataset to which an image or other data file belongs. |
| Resolution | resolution=1 | The image resolution. Each resolution is assigned a key, which is defined in the **res** field of **template_description.json**. |
| Mask | desc="brain", suffix="mask" | Indicates that the image is a binary-valued annotation, where voxels labelled 1 are part of the mask. |
| Discrete segmentation | desc="malf", suffix="dseg" | Indicates that the image is an integer-valued annotation. Each segmentation image file ( **.nii.gz** format) is paired with a dictionary of segment names ( **.tsv** format). |
| Probabilistic segmentation | label="CSF", suffix="probseg" | Indicates that the image is a probabilistic annotation, wherein the value of each voxel indicates the probability of that voxel belonging to the specified label. |
| Atlas | atlas="Schaefer", desc="7Network" | The atlas to which a segmentation file belongs. |
| Transformation | from="MNI152Lin", suffix="xfm" | File containing a mapping between 2 stereotaxic coordinate spaces. The source space is defined in the **from** field, while the target space is defined in the **tpl** field. |
| Image modality | suffix="T1w" | For non-annotation brain images, the suffix indicates whether the image is T1-weighted ( **T1w** ), T2-weighted ( **T2w** ), proton density-weighted ( **PD** ), or T2*-weighted ( **T2star** ). |
| Template cohort | cohort=1 | Subsample of a dataset used to generate an average template. |

**Table S2.** Command-line interface for *TemplateFlow Manager*.

| Argument | Environment variable | Specifications |
| --- | --- | --- |
| `template_id` |  | Identifier of the template. This is the value of the `tpl` field in all file names. |
| `--osf-project` | `OSF_PROJECT` | The OSF project where the template data are to be stored. The project must be writable by the user account whose credentials are specified in the `--osf-user` and `--osf-password` arguments. |
| `--osf-user` | `OSF_USERNAME` | Account username or identifier for OSF cloud storage. |
| `--osf-password` | `OSF_PASSWORD` | Account password for OSF cloud storage. |
| `--osf-overwrite` |  | Flag that indicates that the OSF client should force the overwrite of any existing files in the OSF project that have names conflicting with those of new files. |
| `--gh-user` | `GITHUB_USER` | Account username for GitHub. The user account whose credentials are provided must have a fork of the TemplateFlow repo. |
| `--gh-password` | `GITHUB_PASSWORD` | Account password for GitHub. |
| `--path` | `OSF_PROJECT` | Path to a local directory where template resources are located. The path must either be a directory whose name is `tpl-<template_id>` or contain such a directory. |
| `--nprocs` |  | Maximum number of parallel processes to run when uploading to or fetching from OSF. |

**Table S3.** *TemplateFlow* and NITRC are complementary resources with very different scope, goals and implementations. While *TemplateFlow* is specifically designed for sharing template and atlas resources, the NeuroImaging Tools & Resources Collaboratory (NITRC) “is an award-winning free web-based resource that offers comprehensive information on an ever expanding scope of neuroinformatics software and data” (https://www.nitrc.org/include/about_us.php). Indeed, NITRC provides a fundamental service to the neuroimaging community under open-ended, generalistic purposes, and it has not found a comparable alternative. Indeed, *TemplateFlow* is a narrowly-scoped tool that resolves a very specific set of issues around template, atlases, their sharing and reuse, and their reporting. Reflecting such a hierarchy of resources and acknowledging the relevance of NITRC, *TemplateFlow* was registered therein (https://www.nitrc.org/projects/templateflow).

|  | *TemplateFlow* | NITRC |
| --- | --- | --- |
| Scope | N-D template/atlas resources | General purpose neuroimaging resource cataloging and distribution |
| Storage | Outsourced: leverages (and replicates across) GIN/g-Node, OSF, and Amazon AWS | Fundamental component of NITRC services |
| Data Structure | Data must comply with a BIDS-inspired structure | Fully open to resource owner's choice |
| Programmatic access | Available with *DataLad* and/or the *Python Client* | Not available. Some availability through NITRC-IR, an XNAT instance |
| Ability to store and retrieve individual data files and metadata of templates and atlases | Built-in | Not available |
| Ability to query specific pieces of data or metadata with filters | Yes, based on *PyBIDS* | Limited to Web search or XNAT search when available |
| Versioning | Full version-control with *DataLad* | Limited to creating “snapshot” packages |
| Application Programming Interface (machine operable) | Available, with *DataLad* and/or *PyBIDS* | Not available |
| Quality control | Peer-review at submission time: After submission, feedback is tracked and processed through GitHub Issues | Limited to feedback through Web Discussion Forum, which can be disabled by the owner |
| Unique identifiers | Templates & Sub-template components (e.g., atlases) | Resource |
| Licensing | Resources must be distributed with a license file, which should be permissive and allow derivative works | A license is chosen and can be presented at download time; however, the author is afforded sufficient flexibility to override this |

## Footnotes

1 Author OE notified the authors and the issue is being resolved

2 Author OE reported this practice to the NITRC

## Notes

https://www.templateflow.org/

### Competing Interest Statement

The authors have declared no competing interest.

### Summary of Updates

Revision after a final round of peer-review

## References

Adebimpe A, Bertolero M, Dolui S, Cieslak M, Murtha K, Baller EB, Boeve B, Boxer A, Butler ER, Cook P, Colcombe S, Covitz S, Davatzikos C, Davila DG, Elliott MA, Flounders MW, Franco AR, Gur RE, Gur RC, Jaber B, et al. ASLPrep: A Generalizable Platform for Processing of Arterial Spin Labeled MRI and Quantification of Regional Brain Perfusion. bioRxiv. ; 2021, p. 2021.05.20.444998. doi:10.1101/2021.05.20.444998.

Avants BB, Epstein CL, Grossman M, Gee JC. Symmetric diffeomorphic image registration with cross-correlation: Evaluating auto-mated labeling of elderly and neurodegenerative brain. Med Image Anal. ; 2008, 12(1):26–41. doi:10.1016/j.media.2007.06.004.

Avants BB, Tustison NJ, Song G, Cook PA, Klein A, Gee JC. A reproducible evaluation of ANTs similarity metric performance in brain image registration. NeuroImage. ; 2011, 54(3):2033–44. doi:10.1016/j.neuroimage.2010.09.025.

Blei DM, Ng AY, Jordan MI. Latent Dirichlet Allocation. J Mach Learn Res. ; 2003, 3(Jan):993–1022. https://jmlr.org/papers/v3/blei03a.html.

Bohland JW, Bokil H, Allen CB, Mitra PP. The Brain Atlas Concordance Problem: Quantitative Comparison of Anatomical Parcellations. PLoS One. ; 2009, 4(9):e7200. doi:10.1371/journal.pone.0007200.

Botvinik-Nezer R, Holzmeister F, Camerer CF, Dreber A, Huber J, Johannesson M, Kirchler M, Iwanir R, Mumford JA, Adcock RA, Avesani P, Baczkowski BM, Bajracharya A, Bakst L, Ball S, Barilari M, Bault N, Beaton D, Beitner J, Benoit RG, et al. Variability in the analysis of a single neuroimaging dataset by many teams. Nature. ; 2020, 582(7810):84–88. doi:10.1038/s41586-020-2314-9.

Bowring A, Maumet C, Nichols TE. Exploring the impact of analysis software on task fMRI results. Human Brain Mapping. ; 2019, 40(11):3362–3384. doi:https://doi.org/10.1002/hbm.24603.

Brett M, Johnsrude IS, Owen AM. The problem of functional localization in the human brain. Nat Rev Neurosci. ; 2002, 3:243–249. doi:10.1038/nrn756.

Brodmann K. Brodmann’s: Localisation in the Cerebral Cortex. Springer US; 2006. doi:10.1007/b138298.

Buckner RL, Head D, Parker J, Fotenos AF, Marcus D, Morris JC, Snyder AZ. A unified approach for morphometric and functional data analysis in young, old, and demented adults using automated atlas-based head size normalization: reliability and validation against manual measurement of total intracranial volume. NeuroImage. ; 2004, 23(2):724–738. doi:10.1016/j.neuroimage.2004.06.018.

Calabrese E, Badea A, Watson C, Johnson GA. A quantitative magnetic resonance histology atlas of postnatal rat brain development with regional estimates of growth and variability. NeuroImage. ; 2013, 71:196–206. doi:10.1016/j.neuroimage.2013.01.017.

Carp J. On the Plurality of (Methodological) Worlds: Estimating the Analytic Flexibility of fMRI Experiments. Front Neurosci. ; 2012, 6. doi:10.3389/fnins.2012.00149.

Carp J. The secret lives of experiments: Methods reporting in the fMRI literature. NeuroImage. ; 2012, 63(1):289–300. doi:10.1016/j.neuroimage.2012.07.004.

Coalson TS, Van Essen DC, Glasser MF. The impact of traditional neuroimaging methods on the spatial localization of cortical areas. Proceedings of the National Academy of Sciences. ; 2018, 115(27):E6356–E6365. doi:10.1073/pnas.1801582115.

Collins DL, Neelin P, Peters TM, Evans AC. Automatic 3D Intersubject Registration of MR Volumetric Data in Standardized Talairach Space. Journal of Computer Assisted Tomography. ; 1994, 18(2):192–205. https://insights.ovid.com/crossref?an=00004728-199403000-00005.

Courchesne E, Chisum HJ, Townsend J, Cowles A, Covington J, Egaas B, Harwood M, Hinds S, Press GA. Normal Brain Development and Aging: Quantitative Analysis at in Vivo MR Imaging in Healthy Volunteers. Radiology. ; 2000, 216(3):672–682. doi:10.1148/radiology.216.3.r00au37672.

Cox RW, Hyde JS. Software tools for analysis and visualization of fMRI data. NMR Biomed. ; 1997, 10(4-5):171–178. doi:10.1002/(SICI)1099-1492(199706/08)10:4/5<171::AID-NBM453>3.0.CO;2-L.

Dickie DA, Job DE, Gonzalez DR, Shenkin SD, Wardlaw JM. Use of Brain MRI Atlases to Determine Boundaries of Age-Related Pathology: The Importance of Statistical Method. PLoS One. ; 2015, 10(5):e0127939. doi:10.1371/journal.pone.0127939.

Dickie DA, Shenkin SD, Anblagan D, Lee J, Blesa Cabez M, Rodriguez D, Boardman JP, Waldman A, Job DE, Wardlaw JM. Whole Brain Magnetic Resonance Image Atlases: A Systematic Review of Existing Atlases and Caveats for Use in Population Imaging. Front Neuroinform. ; 2017, 11. doi:10.3389/fninf.2017.00001.

Esteban O, Birman D, Schaer M, Koyejo OO, Poldrack RA, Gorgolewski KJ. MRIQC: Advancing the Automatic Prediction of Image Quality in MRI from Unseen Sites. PLoS One. ; 2017, 12(9):e0184661. doi:10.1371/journal.pone.0184661.

Esteban O, Markiewicz CJ, Blair RW, Moodie CA, Isik AI, Erramuzpe A, Kent JD, Goncalves M, DuPre E, Snyder M, Oya H, Ghosh SS, Wright J, Durnez J, Poldrack RA, Gorgolewski KJ. fMRIPrep: a robust preprocessing pipeline for functional MRI. Nat Meth. ; 2019, 16(1):111–116. doi:10.1038/s41592-018-0235-4.

Evans AC, Collins DL, Mills SR, Brown ED, Kelly RL, Peters TM. 3D statistical neuroanatomical models from 305 MRI volumes. In: IEEE Conference Record Nuclear Science Symposium and Medical Imaging Conference, vol. 3 San Francisco, CA, USA; 1993. p. 1813–1817. doi:10.1109/NSSMIC.1993.373602.

Evans AC, Janke AL, Collins DL, Baillet S. Brain templates and atlases. NeuroImage. ; 2012, 62(2):911–922. doi:10.1016/j.neuroimage.2012.01.024.

Fischl B, Sereno MI, Dale AM. Cortical surface-based analysis II: Inflation, flattening, and a surface-based coordinate system. NeuroImage. ; 1999, 9(2):195–207.

Fonov V, Evans AC, Botteron K, Almli CR, McKinstry RC, Collins DL. Unbiased average age-appropriate atlases for pediatric studies. NeuroImage. ; 2011, 54(1):313–327. doi:10.1016/j.neuroimage.2010.07.033.

Fonov V, Evans A, McKinstry R, Almli C, Collins D. Unbiased nonlinear average age-appropriate brain templates from birth to adulthood. NeuroImage. ; 2009, 47, Supplement 1:S102. doi:10.1016/S1053-8119(09)70884-5.

Frey S, Pandya DN, Chakravarty MM, Petrides M, Collins DL. MNI monkey space. Neuroscience Research. ; 2009, 65:S130. doi:10.1016/j.neures.2009.09.629.

Friston KJ, Ashburner J, Kiebel SJ, Nichols TE, Penny WD. Statistical parametric mapping : the analysis of functional brain images. London: Academic Press; 2006.

Goerzen D, Fowler C, Devenyi GA, Germann J, Madularu D, Chakravarty MM, Near J. An MRI-Derived Neuroanatomical Atlas of the Fischer 344 Rat Brain. Sci Rep. ; 2020, 10(1):6952. doi:10.1038/s41598-020-63965-x.

Good CD, Johnsrude IS, Ashburner J, Henson RN, Friston KJ, Frackowiak RS. A voxel-based morphometric study of ageing in 465 normal adult human brains. NeuroImage. ; 2001, 14(1):21–36. doi:10.1006/nimg.2001.0786.

Gorgolewski KJ, Auer T, Calhoun VD, Craddock RC, Das S, Duff EP, Flandin G, Ghosh SS, Glatard T, Halchenko YO, Handwerker DA, Hanke M, Keator D, Li X, Michael Z, Maumet C, Nichols BN, Nichols TE, Pellman J, Poline JB, et al. The brain imaging data structure, a format for organizing and describing outputs of neuroimaging experiments. Sci Data. ; 2016, 3:160044. doi:10.1038/sdata.2016.44.

Grabner G, Janke AL, Budge MM, Smith D, Pruessner J, Collins DL. Symmetric Atlasing and Model Based Segmentation: An Application to the Hippocampus in Older Adults. In: Larsen R, Nielsen M, Sporring J, editors. Medical Image Computing and Computer-Assisted Intervention – MICCAI 2006 Lecture Notes in Computer Science, Berlin, Heidelberg: Springer; 2006. p. 58–66. doi:10.1007/11866763_8.

Halchenko YO. Incorrect probabilities in Harvard-Oxford-sub Left hemisphere. FSL–users mailing list. ; 2013. https://www.jiscmail.ac.uk/cgi-bin/webadmin?A2=FSL;2bb44bee.1301.

Halchenko YO, Meyer K, Poldrack B, Solanky DS, Wagner AS, Gors J, MacFarlane D, Pustina D, Sochat V, Ghosh SS, Mönch C, Markiewicz CJ, Waite L, Shlyakhter I, Vega Adl, Hayashi S, Häusler CO, Poline JB, Kadelka T, Skytén K, et al. DataLad: distributed system for joint management of code, data, and their relationship. Journal of Open Source Software. ; 2021, 6(63):3262. doi:10.21105/joss.03262.

Holmes CJ, Hoge R, Collins L, Woods R, Toga AW, Evans AC. Enhancement of MR Images Using Registration for Signal Averaging. J Comput Assist Tomogr. ; 1998, 22(2):324–333. https://insights.ovid.com/crossref?an=00004728-199803000-00032.

Jung B, Taylor PA, Seidlitz J, Sponheim C, Perkins P, Ungerleider LG, Glen D, Messinger A. A comprehensive macaque fMRI pipeline and hierarchical atlas. NeuroImage. ; 2021, 235:117997. doi:10.1016/j.neuroimage.2021.117997.

Kazhdan M, Hoppe H. Screened Poisson Surface Reconstruction. ACM Transactions on Graphics. ; 2013, 32(3):29:1–29:13. doi:10.1145/2487228.2487237.

Landman BA, Ribbens A, Lucas B, Davatzikos C, Avants B, Ledig C, Ma D, Rueckert D, Vandermeulen D, Maes F. MICCAI 2012 Workshop on Multi-Atlas Labeling. CreateSpace Independent Publishing Platform; 2012.

Lüders E, Steinmetz H, Jäncke L. Brain size and grey matter volume in the healthy human brain. NeuroReport. ; 2002, 13(17):2371–2374. https://journals.lww.com/neuroreport/Abstract/2002/12030/Brain_size_and_grey_matter_volume_in_the_healthy.40.aspx.

MacNicol E, Ciric R, Kim E, Censo DD, Cash D, Poldrack R, Esteban O. Atlas-based brain extraction is robust across rat MRI studies. In: IEEE 19th International Symposium on Biomedical Imaging (ISBI 2021) Nice, France; 2021. p. (accepted). doi:10.1109/ISBI48211.2021.9.

MacNicol E, Wright P, Kim E, Brusini I, Esteban O, Simmons C, Turkheimer F, Cash D. Age-specific adult rat brain MRI templates and tissue probability maps. Frontiers in Neuroinformatics. ; 2021. doi:10.3389/fninf.2021.669049.

Markiewicz CJ, Gorgolewski KJ, Feingold F, Blair R, Halchenko YO, Miller E, Hardcastle N, Wexler J, Esteban O, Goncavles M, Jwa A, Poldrack R. The OpenNeuro resource for sharing of neuroscience data. eLife. ; 2021, 10:e71774. doi:10.7554/eLife.71774.

Mars RB, Passingham RE, Jbabdi S. Connectivity Fingerprints: From Areal Descriptions to Abstract Spaces. Trends Cogn Sci. ; 2018, 22(11):1026–1037. doi:10.1016/j.tics.2018.08.009.

Martin RF, Bowden DM. Primate Brain Maps: Structure of the Macaque Brain. Elsevier; 2000.

Matsuzawa J, Matsui M, Konishi T, Noguchi K, Gur RC, Bilker W, Miyawaki T. Age-related Volumetric Changes of Brain Gray and White Matter in Healthy Infants and Children. Cereb Cortex. ; 2001, 11(4):335–342. doi:10.1093/cercor/11.4.335.

Mazziotta J, Toga A, Evans A, Fox P, Lancaster J, Zilles K, Woods R, Paus T, Simpson G, Pike B, Holmes C, Collins L, Thompson P, MacDonald D, Iacoboni M, Schormann T, Amunts K, Palomero-Gallagher N, Geyer S, Parsons L, et al. A Four-Dimensional Probabilistic Atlas of the Human Brain. J Am Med Inform Assoc. ; 2001, 8(5):401–430. doi:10.1136/jamia.2001.0080401.

Mazziotta JC, Toga AW, Evans A, Fox P, Lancaster J. A Probabilistic Atlas of the Human Brain: Theory and Rationale for Its Development: The International Consortium for Brain Mapping (ICBM). NeuroImage. ; 1995, 2(2, Part A):89–101. doi:10.1006/nimg.1995.1012.

Papp EA, Leergaard TB, Calabrese E, Johnson GA, Bjaalie JG. Waxholm Space atlas of the Sprague Dawley rat brain. NeuroImage. ; 2014, 97:374–386. doi:10.1016/j.neuroimage.2014.04.001.

Paxinos G, Watson C. The rat brain in stereotaxic coordinates. Elsevier Academic Press; 1997.

Pisner D, Hammonds R. PyNets: A Reproducible Workflow for Structural and Functional Connectome Ensemble Learning. In: Annual Meeting of the Organization for Human Brain Mapping, vol. 26 Online Event; 2020. p. 1967. https://github.com/dPys/PyNets/.

Robinson EC, Jbabdi S, Glasser MF, Andersson J, Burgess GC, Harms MP, Smith SM, Van Essen DC, Jenkinson M. MSM: A new flexible framework for Multimodal Surface Matching. NeuroImage. ; 2014, 100:414–426. doi:10.1016/j.neuroim-age.2014.05.069.

Rohlfing T. Incorrect ICBM-DTI-81 atlas orientation and white matter labels. Front Neurosci. ; 2013, 7. doi:10.3389/fnins.2013.00004.

Satterthwaite TD, Connolly JJ, Ruparel K, Calkins ME, Jackson C, Elliott MA, Roalf DR, Hopson R, Prabhakaran K, Behr M, Qiu H, Mentch FD, Chiavacci R, Sleiman PMA, Gur RC, Hakonarson H, Gur RE. The Philadelphia Neurodevelopmental Cohort: A publicly available resource for the study of normal and abnormal brain development in youth. NeuroImage. ; 2016, 124:1115–1119. doi:10.1016/j.neuroimage.2015.03.056.

Sawiak SJ, Wood NI, Williams GB, Morton AJ, Carpenter TA. SPMMouse: A new toolbox for SPM in the animal brain. In: Proc. Intl. Soc. Mag. Reson. Med., vol. 17 Hawaii, USA; 2009. p. 1086.

Schilling KG, Gao Y, Stepniewska I, Wu TL, Wang F, Landman BA, Gore JC, Chen LM, Anderson AW. The VALiDATe29 MRI Based Multi-Channel Atlas of the Squirrel Monkey Brain. Neuroinform. ; 2017, 15(4):321–331. doi:10.1007/s12021-017-9334-0.

Schurr PH, Merrington WR. The Horsley–Clarke stereotaxic apparatus. Br J Surg. ; 1978, 65(1):33–36. doi:10.1002/bjs.1800650110.

Shi F, Yap PT, Wu G, Jia H, Gilmore JH, Lin W, Shen D. Infant Brain Atlases from Neonates to 1- and 2-Year-Olds. PLoS One. ; 2011, 6(4):1–11. doi:10.1371/journal.pone.0018746.

Sowell ER, Peterson BS, Thompson PM, Welcome SE, Henkenius AL, Toga AW. Mapping cortical change across the human life span. Nat Neurosci. ; 2003, 6(3):309–315. doi:10.1038/nn1008.

Szulc KU, Lerch JP, Nieman BJ, Bartelle BB, Friedel M, Suero-Abreu GA, Watson C, Joyner AL, Turnbull DH. 4D MEMRI atlas of neonatal FVB/N mouse brain development. NeuroImage. ; 2015, 118:49–62. doi:10.1016/j.neuroimage.2015.05.029.

Talairach J, Tournoux P. Co-planar stereotaxic atlas of the human brain. Stuttgart New York: Georg Thieme Verlag/Thieme Medical Publishers; 1988.

Talairach J, David M, Tournoux P, Corredor H, Kvasina J. Atlas d’anatomie stéréotaxique: repérage radiologique indirect des noyaux gris centraux des régions mésencéphalo-sous-optique et hypothalamique de l’homme. Masson; 1957.

Tavor I, Jones OP, Mars RB, Smith SM, Behrens TE, Jbabdi S. Task-free MRI predicts individual differences in brain activity during task performance. Science. ; 2016, 352(6282):216–220. doi:10.1126/science.aad8127.

Thompson W, Fanton S. Open Source Software: NetPlotBrain. Zenodo. ; 2021. doi:10.5281/zenodo.4593837.

Tosun D, Siddarth P, Levitt J, Caplan R. Cortical thickness and sulcal depth: insights on development and psychopathology in paediatric epilepsy. BJPsych Open. ; 2015, 1(2):129–135. doi:10.1192/bjpo.bp.115.001719.

Tustison NJ, Avants BB, Cook PA, Zheng Y, Egan A, Yushkevich PA, Gee JC. N4ITK: Improved N3 Bias Correction. IEEE Transactions on Medical Imaging. ; 2010, 29(6):1310–1320. doi:10.1109/TMI.2010.2046908.

Van Essen DC. Windows on the brain: the emerging role of atlases and databases in neuroscience. Curr Opin Neurobiol. ; 2002, 12(5):574–579. doi:10.1016/S0959-4388(02)00361-6.

Van Essen DC, Glasser MF, Dierker DL, Harwell J, Coalson T. Parcellations and Hemispheric Asymmetries of Human Cerebral Cortex Analyzed on Surface-Based Atlases. Cerebral Cortex. ; 2012, 22(10):2241–2262. doi:10.1093/cercor/bhr291.

Von Economo CF, Koskinas GN. Atlas of cytoarchitectonics of the adult human cerebral cortex, vol. 10. Karger Basel; 2008.

Wang H, Suh JW, Das SR, Pluta JB, Craige C, Yushkevich PA. Multi-Atlas Segmentation with Joint Label Fusion. IEEE Transactions on Pattern Analysis and Machine Intelligence. ; 2013, 35(3):611–623. doi:10.1109/TPAMI.2012.143.

Wilkinson MD, Dumontier M, Aalbersberg IJ, Appleton G, Axton M, Baak A, Blomberg N, Boiten JW, da Silva Santos LB, Bourne PE, Bouwman J, Brookes AJ, Clark T, Crosas M, Dillo I, Dumon O, Edmunds S, Evelo CT, Finkers R, Gonzalez-Beltran A, et al. The FAIR Guiding Principles for scientific data management and stewardship. Sci Data. ; 2016, 3(1):160018. doi:10.1038/sdata.2016.18.

Yarkoni T, Markiewicz C, de la Vega A, Gorgolewski K, Salo T, Halchenko Y, McNamara Q, DeStasio K, Poline JB, Petrov D, Hayot-Sasson V, Nielson D, Carlin J, Kiar G, Whitaker K, DuPre E, Wagner A, Tirrell L, Jas M, Hanke M, et al. PyBIDS: Python tools for BIDS datasets. J Open Source Softw. ; 2019, 4:1294. doi:10.21105/joss.01294.

Yeo BT, Krienen FM, Sepulcre J, Sabuncu MR, Lashkari D, Hollinshead M, Roffman JL, Smoller JW, Zöllei L, Polimeni JR, Fischl B, Liu H, Buckner RL. The organization of the human cerebral cortex estimated by intrinsic functional connectivity. J Neurophysiol. ; 2011, 106(3):1125–1165. doi:10.1152/jn.00338.2011.

Yoon U, Fonov VS, Perusse D, Evans AC. The effect of template choice on morphometric analysis of pediatric brain data. NeuroImage. ; 2009, 45(3):769–777. doi:10.1016/j.neuroimage.2008.12.046.

